# Persistent ECM Scarring Reprograms Intestinal Stem Cells to Drive Chronic Inflammation

**DOI:** 10.1101/2025.06.08.658481

**Authors:** Idan Adir, Carmel Sochen, Aviya Habshush Menachem, Sacha Lebon, Barak Toval, Vladyslav Holiar, Natalia Davidzohn, Tomer-Meir Salame, Irit Rosenhek-Goldian, Simonas Savickas, Fabio Sabino, Inna Solomonov, Ulrich Auf dem Keller, Moshe Biton, Irit Sagi

## Abstract

Tissue regeneration is conventionally viewed as a return to homeostasis. Here, we uncover that the extracellular matrix (ECM) in the colon undergoes a lasting pathological reprogramming following inflammation, forming a remodeled niche—modECM—that persistently disrupts intestinal stem cell (ISC) identity. Using temporal multi-omics, biomechanical profiling, and spatial fate mapping in murine colitis models, we show that modECM, characterized by Collagen XVIII accumulation and immune-driven proteolysis, redirects ISCs toward a wound-associated, squamous-like epithelial state with pro-inflammatory transcriptional signatures. Ex vivo, modECM alone reprograms ISC fate by suppressing Wnt signaling and activating immune recruitment pathways. In vivo, modECM-rich zones sustain T cell infiltration and KRT14⁺ epithelial cell emergence from Lgr5⁺ progenitors. This aberrant epithelial program is mirrored in inflamed rectal biopsies from ulcerative colitis patients. Our findings redefine the ECM as a long-lived instructive compartment that encodes injury memory and promotes maladaptive regeneration, positioning it as a potential therapeutic target in chronic inflammatory disease.

## Introduction

Gut pathologies, including inflammatory bowel disease (IBD), emerge from a complex interplay between tissue damage, inflammation, and the regenerative capacity of intestinal stem cells (ISCs). The ability of ISCs to regenerate the intestinal epithelium is critical for restoring barrier integrity and maintaining homeostasis; however, this process can be profoundly influenced by the surrounding microenvironment^1–3^. To understand how ISCs adapt to different physiological and pathological conditions, it is essential to closely examine their local environment, known as the stem cell niche^4–6^. The ISC niche consists of a diverse array of cell types, including epithelial, stromal, such as crypt fibroblasts and endothelial cells, as well as neurons and both innate and adaptive immune cells. These components are spatially organized to meet the unique needs of the tissue, allowing the stem cells to respond effectively to a complex network of cellular and extracellular signals^2,5–10^. ISC function is also regulated by the extracellular matrix (ECM), which provides structural and mechanical support as well as biochemical cues that guide stem cell behavior^11–13^. Yet, efforts to decipher the mechanisms underlying tissue repair and regeneration have focused largely on the cellular processes of stem cell renewal, differentiation, proliferation, and their interactions with immune and stromal cells^14–19^, while the ECM-ISC cross-talk in the context of post-inflammation regeneration and tissue damage remains largely understudied. Specifically, it is unclear how widespread insults, such as infection or chronic inflammation, which are known to affect the mucosal ECM, are transmitted to the tissue-resident stem cell pool. Previous studies have highlighted the crucial role of ECM structural features and its matrisome, comprising proteins and affiliated factors involved in ECM remodeling, in gut inflammation^20–22^. However, its role as a reservoir of signaling molecules capable of serving as stem cell modulators in tissue regeneration and recovery is underappreciated.

In the inflammatory phase of colitis, innate immune cells, in particular neutrophils and monocytes, are recruited to the inflammatory site and secrete ECM-remodeling enzymes such as matrix-metalloproteinases (MMPs and ADAMs)^23–25^. These innate immune cells elicit an inflammatory response to promote tissue regeneration by repair factors such as fibroblast growth factors (FGFs) and transforming growth factor beta (TGF-β), respectively. FGF and TGFΒ accumulation result in fibroblast recruitment to the inflammatory site, depositing ECM components, including fibronectin, hyaluronic acid, proteoglycans, and different collagens^21,26^. However, in chronic gut inflammation, the fibroblast state may shift to inflammatory-associated fibroblasts (IAFs), resulting in promoting the inflammatory state of the tissue by depositing pro-inflammatory factors such as Collagen XVIII (*Col18a1*) and interleukin 11 (*IL-11*)^13,27^. These IAF subsets were recently shown to persist longer after the inflammatory phase but are cleared from the tissue during the recovery state^7^. In IBD mouse models and patients, structural changes in the ECM are noticeable even before the onset of inflammation^20,22,28^. However, whether the ECM reverts to normal or remains damaged in the recovery phase, and how it might shape the regenerative landscape in chronic gut inflammation, has not been assessed.

At barrier surfaces, such as the intestine, epithelial cells also play a critical role in tissue regeneration by undergoing a dynamic transition into wound-associated epithelial (WAE) cells to restore barrier integrity in response to inflammation^29,30^. At the regenerative phase, WAE cells depolarize, proliferate, and migrate to the damaged permeable tissue sites to restore barrier integrity by epithelial restitution. Despite the recognized importance of WAE cells in epithelial restitution, it remains unclear how the WAE cell transition occurs. A recent study defined the WAE cell differentiation program at single cell resolution and demonstrated that WAE cells instruct fibroblasts to promote tissue healing through ECM remodeling^31^. Indeed, a recent study showed that the ECM factor, laminin in the basement membrane, is essential for maintaining Lgr5+ ISCs and promoting crypt-like morphology^32^. However, how the recovered ECM state affects intestinal stem cell (ISC) regeneration is still unexplored.

Here, we show that the mouse mucosa’s ECM remained altered even after the recovery phase in the case of experimental colitis, due to the proteolytic activity of myeloid cells. This leads to basal membrane disruption, decreased ECM stiffness, and an increase in ECM IBD biomarkers, e.g., collagen XVIII. Through a combination of proteomic, biomechanical, and imaging analyses, we also identified distinct changes in the protein composition of the ECM critical for maintaining the ISC niche, such as a reduction in Wnt and an elevation of BMP ligands, respectively. Changes in cryptal ECM led to the conversion of Lgr5+ ISC to WAE-like cells with a distinct transcriptional program associated with T cell recruitment signals. Thus, the modified ECM, which we refer to as modECM, mediates pathophysiological cellular transitions that have a lasting detrimental effect on tissue regeneration, potentially perpetuating chronic inflammatory conditions. These findings may implicate the ECM’s regulatory role in other tissues and suggest that abnormal reprogramming into modECM is likely involved in other inflammatory-related pathologies.

## Results

### Acute experimental colitis results in persistent inflammation and abnormal tissue recovery

To investigate the role of the ECM in the regeneration of the colon tissue after inflammation, we utilized the well-established mouse model of dextran sulfate sodium (DSS)-induced acute colitis^33^ (**Figure 1A**). DSS increased permeability of the epithelial barrier, resulting in an immune response and subsequent ulceration. The DSS-induced colitis model is characterized by an acute phase (Day 10), marked by epithelial barrier disruption, inflammation, and weight loss, followed by a regenerative phase (Day 21), during which tissue repair, crypt remodeling, and gradual restoration of epithelial integrity occur ^34^. Mice were given a low dose of DSS (1.25%) in their drinking water for 7 days and were then allowed to recover for the remainder of the experiment. To identify key regeneration time points, we monitored clinical manifestations, including weight loss, a hallmark of acute DSS-induced colitis. Mice began losing weight on days 6–7, with a maximum loss of <20% by days 10–15. Recovery was rapid, and by Day 38, body mass returned to baseline, eventually following a growth pattern similar to healthy mice. (**Figure 1B**). Despite the increase in animal weight at the recovery phase, clinical signs of colitis could still be observed in the colonoscopies of mice^35^ as far as Day 80 post-DSS administration (**Figures 1C, S1A-B**). Most mice presented with inflamed colons (52% inflamed and 4% severely inflamed) by Day 10. Pathological signs included opaque colon walls, fibrin deposits, and soft feces. By Day 38, colon inflammation decreased to 26.7%, with reduced fibrin deposits, granularity, and feces softness, despite minor bleeding. By Day 80, inflammation persisted, and colon width remained enlarged as in the acute phase (**Figures S1C-D**). We assessed colonic epithelial barrier regeneration using a gut permeability assay with dextran-FITC at different time points after DSS induction. The results showed a significant barrier breach during the acute phase (Day 10, ∼8-fold increase vs. healthy colons), which gradually improved, returning to normal by Day 80 (**Figure 1D**). Histological staining (H&E and Mason Trichrome (MT), **Figure 1E**) revealed ulcerations and epithelial loss at the acute phase (Day 10), along with tertiary lymphoid structures (TLS) surrounding crypts and ulcers (**Figure 1F**). TLS persisted through Days 38 and 80. Remarkably, during the recovery phase, colons displayed abnormal mucosal regions, including branched crypts associated with fibrosis, suggesting abnormal regeneration of the mucosal ECM (**Figure 1E**). The recovery time points were also characterized by squamous-like epithelium in the far distal colon. This phenomenon has been recently described as the migration of neighboring non-colonic skin-like cells from the anorectal transitional zone (TZ) to squamous neo-epithelial cells (SNECs) ^36,37^. Of Note, the aberration in tissue histology persisted up to Day 400 in this model **(Figure S1E)**. The observed abnormal recovery of the mucosal ECM and invasion of SNECs from the distal rectum prompted us to further investigate the molecular mechanisms underlying the modulation of the mucosal ECM and its influence on Lgr5+ ISCs during the regeneration and recovery phases of colon glandular epithelium.

**Figure 1.**
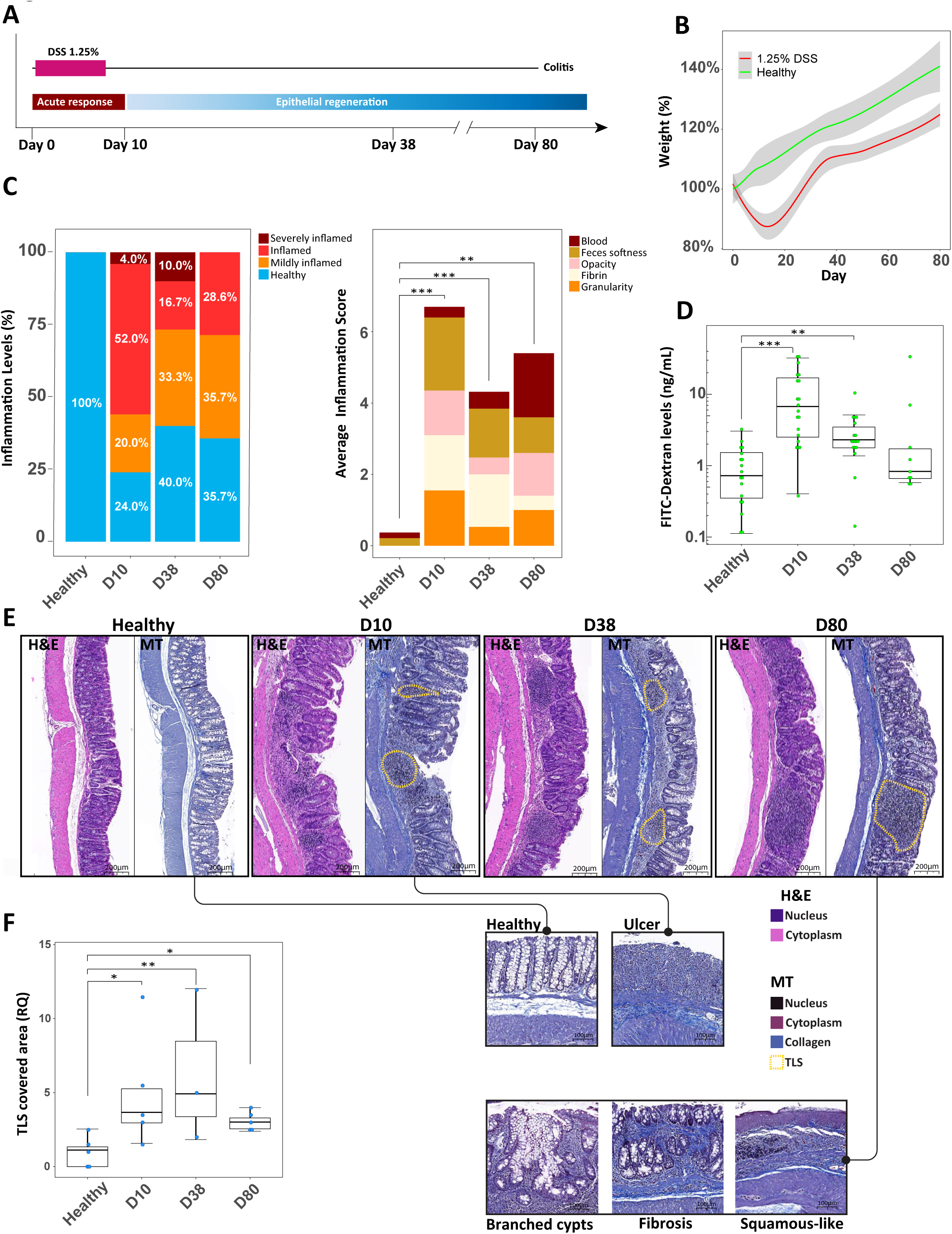
Abnormal recovery of the colon in the DSS-induced colitis model. **(A)** Scheme of DSS-induced colitis experiment. (**B**) Change in body weight of mice after DSS induction. Weight progression of healthy mice (green) or DSS-induced colitis mice (red), standard error in grey. (**C**) Left-Stacked bar graph of inflammation score percentage per timepoint, legends on top right. Right-Colonoscopy scores were converted to inflammation level categories. *\*\* p≤0.01, *** p≤0.001 one way ANOVA, n ≥* 5 mice per group, legends on top right. (**D**) Permeability assay of DSS timepoints. Values are the relative concentration (ng/mL) of dextran FITC in the serum of mice compared to the healthy mice average. *\*\* p≤0.01, *** p≤0.001 log transformed one way ANOVA, n ≥* 9 mice per group. (**E**) Hematoxylin and Eosin (H&E) and Masson’s trichrome (MT) staining of colon sections from the DSS timeline, tissue lymphoid structures (TLSs) marked with a yellow outline, scale bar of 200μm. Bottom-Representative images of normal (top left) or abnormal mucosal morphologies, such as ulcers in Day 10 (top right) and ones found in Day 80 tissue (bottom panels), scale bar of 100μm. (**F**) Relative quantification of the TLS-covered area out of the colon section (percentage) compared to the healthy mean. *\* p≤0.05, ** p≤0.01, log transformed one way ANOVA, n ≥* 3 mice per group.

### Infiltrating myeloid immune cells secreting ECM-degrading enzymes during gut inflammation

To further characterize the inflammatory response of DSS colitis, we used single-cell CyTOF analysis to map the immune cell composition and ECM remodeling enzyme repertoire in injured colon tissues (**Figure 2**). Immune cell populations varied across time points, showing a shift from myeloid to lymphoid lineages from Day 10 through Day 38 to Day 80. **(Figures 2A, S2)**. The levels of infiltrating myeloid cell lineages, including macrophages, neutrophils, and dendritic cells, peaked on the acute inflammatory phase of Day 10, and then gradually decreased **(Figure 2B)**. Conversely, the levels of infiltrating lymphoid lineage cells, including B, T, and NK cells, gradually increased throughout the recovery phase, peaking at Day 80 **(Figure 2C)**. Of note, T cells were elevated about 5-fold more compared to healthy tissue on Day 80. T cells comprise both effectors and regulatory T helper cells that are known to play a major role in the impaired immune tolerance known to occur in IBD ^38,39^. Our CyTOF results uncovered that MMP9, a known protease of collagen type IV ^40,41^, was highly expressed by neutrophils and iNOS^+^ macrophages at the inflammation peak (Day 10), and in Day 38 of tissue recovery by neutrophils (**Figures 2D-E**). Accordingly, collagen type IV was elevated in our degradome data (**Methods** and **Figure 3**), suggesting the involvement of neutrophils and iNOS⁺ macrophages in Collagen IV degradation. Another ECM remodeling enzyme implicated in inflammation was produced by specific cell types. MMP14, a membrane-bound collagenase, was exclusively expressed by iNOS⁺ macrophages, whereas MMP2, a constitutively expressed gelatinase requiring activation by other MMPs, was produced by all cell types at different time points. These findings indicate that during peak inflammation, ECM degradation or its damage is primarily driven by iNOS⁺ macrophages, while by Day 38, during the “recovery” phase, neutrophils contribute to basement membrane degradation.

**Figure 2.**
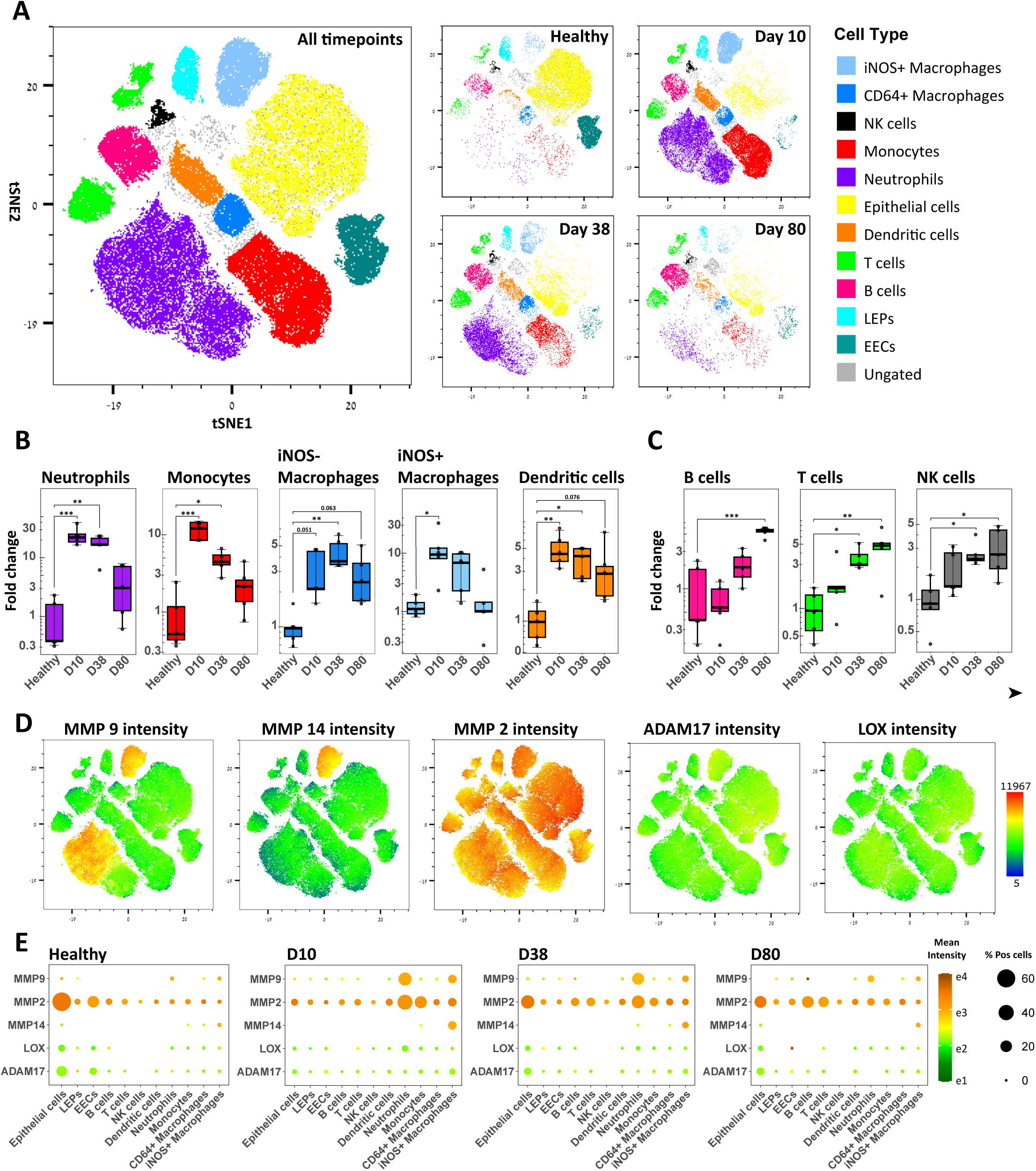
Immune cells analysis in the colon of DSS-treated mice. (**A**) Left – t-SNE plot made from cytometry by time of flight (cyTOF) analysis of all DSS-timepoints. Right - separation of the t-SNE into the four representative timepoints: Healthy, Days 10, 38, and 80. Cells in the t-SNE plot were separated into spatially distinct subsets and identified according to a combination of markers, see **Figure S2**. Legends, far right. (**B,C**) Myeloid (B) or Lymphoid (C) cell populations measured (percentage) out of all live cells. Cell type on top of each plot. *\* p≤0.05, ** p≤0.01, *** p≤0.001, one way ANOVA. n =* 5 mice per group. (**D**) ECM remodeling enzyme mean intensity displayed color in the tSNE plot. Legends, far right. (**E**) ECM remodeling enzyme expression per cell type per time point. The color of dots indicates mean expression intensity, and the size of dots indicates the percentage of positive cells for the remodeling enzyme out of all cells.

**Figure 3.**
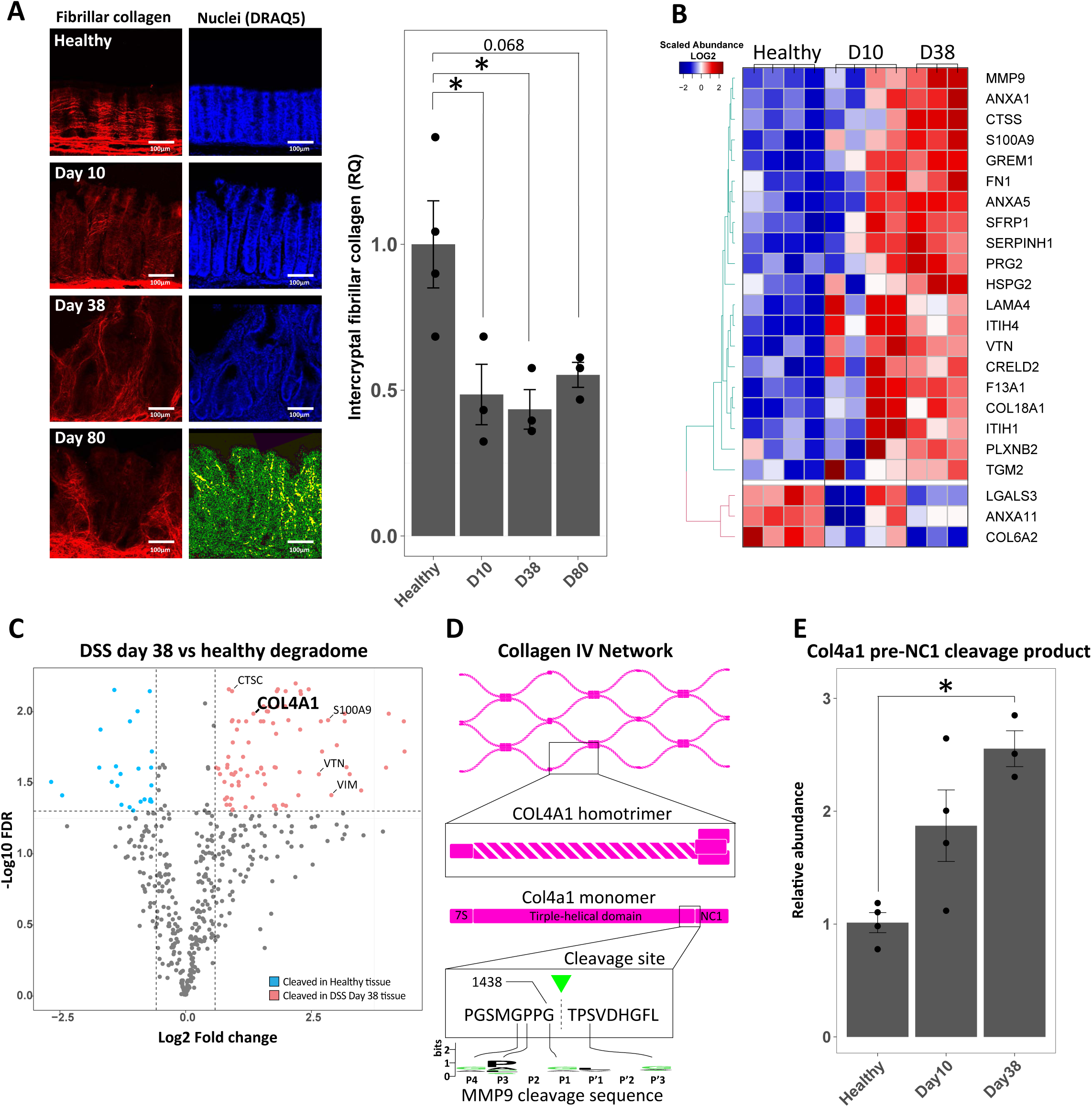
Mucosal ECM alteration in DSS timepoints. (**A**) Second harmonic generation (SHG) imaging of intercryptal collagen. Left, representative images of Fibrillar collagen (red) and Nuclei (blue) from healthy, acute (Day 10), and regenerated (Day 38 and 80) in DSS time points, scale bar of 100μm. Right, relative quantification of intercryptal fibrillar collagen compared to healthy, units are average intensity per crypt per mouse, *\* p≤0.05, log transformed one way ANOVA, n ≥* 3 mice per group (4 regions per mouse). (**B**) Heatmap of differentially expressed matrisome proteins from whole colon tissue of healthy, Days 10 and 38 of DSS-treated mice measured by LC-MS/MS. Color legends of Log2 expression, top left. (**C**) Volcano plot of differentially expressed neo-N-termini degradome products, comparing DSS Day 38 to healthy tissue, *n ≥ 3* mice per group. Legends are on the far right of the plot. (**D**) Top-schematic of COL4A1 neo-N-terminus peptide location found in degradomic analysis. Bottom-comparison between cleavage site sequence in COL4A1 and MMP9 preferred cleavage sequence as found in MEROPS database. (**E**) Relative abundance of COL4A1 neo-N-terminus peptides compared to healthy mice. *\* p≤0.05, FDR post-hoc of student’s t-test, n ≥ 3* mice per group.

### Mucosal ECM damage persists beyond tissue recovery

The immune cell infiltration and mucosal morphological changes persisted throughout the recovery period and beyond, lasting up to 400 days, as shown by the MT images (**Figure S1E**). These findings suggest long-lasting damage to the mucosal ECM, affecting both its structure and composition. Using second harmonic generation two-photon imaging, we detected a significant decrease in collagen fibers on Days 38 and 80, indicating intercryptal fibrillar collagen degradation and long-lasting ECM damage. The observed aberrant ECM alterations directly impacted the abnormal formation of cryptal-cell morphologies, as shown by DAPI staining (**Figure 3A**), aligning with the branched crypt morphologies observed in histology (**Figure 1E**). To quantify potential alterations in ECM protein composition, we performed liquid chromatography-tandem mass spectrometry (LC-MS/MS) proteomics on tissue samples collected on Days 10 and 38 after DSS administration. Among the 1,832 identified proteins (**Table S1**), 94 were of matrisome or matrisome-associated proteins. We observed a significant increase in several structural ECM components, including Collagen XVIII (COL18A1), previously identified as an early IBD marker^20^, Fibronectin 1 (FN1), Vitronectin (VTN), and Matrix Metalloproteinase 9 (MMP9) (**Figure 3B**). To further characterize the activity of ECM-related proteases, we performed degradomic analysis of colon tissues (**Methods**) from Day 0 to 38 to quantify changes in the abundance of neo-N-termini cleavage products^42^. Among the 477 neo-N-termini detected across different time points, we identified a significant increase in a single cleavage site within collagen IV subunit A1 (COL4A1), a basement membrane collagen, occurring just before its first non-collagenous (NC1) domain (**Figures 3C-E, Table S2**). This specific COL4A1 cleavage site has been previously described as an MMP9 substrate ^43^, Proline at P3 and Glycine at P4 and P1^44^ (**Figure 3D**). These results support the observation of abnormal accumulation of COL18A1 and degradation of COL4A1 during the recovery phase at the tissue level (**Figures 3B,C**, and **Table S1**). Interestingly, COL18A1 was shown recently as a marker of a subset of inflammatory-associated fibroblasts-3 (IAF3), found to be enriched in gut inflammation ^27^ but reduced in the recovery phase^7^. We validated the expansion of IAF-3 in the inflammatory phase (Day 10) and their decline in the recovery phase (days 38 and 80), using immunofluorescence assay (IFA) of COL18A1, PDPN and VCAM1, markers of IAF3, as reported before^7^ in the pericryptal regions (**Figures 4A,B**), as well as in the ulcers of the acute inflammation at Day 10 (**Figure S3A**), and expanded lamina propria and squamous-like epithelium of the recovery timepoints Days 38 and 80 (**Figure S3B-C**). However, the level of COL18A1 in the tissue remained high along the examined days (**Figure 4C**), especially in the distal part of the colon, indicating that COL18A1 is sequestered by the ECM and remains embedded in the deformed matrix long after tissue regeneration and recovery phases, suggesting of lasting ECM modification after gut inflammation.

**Figure 4.**
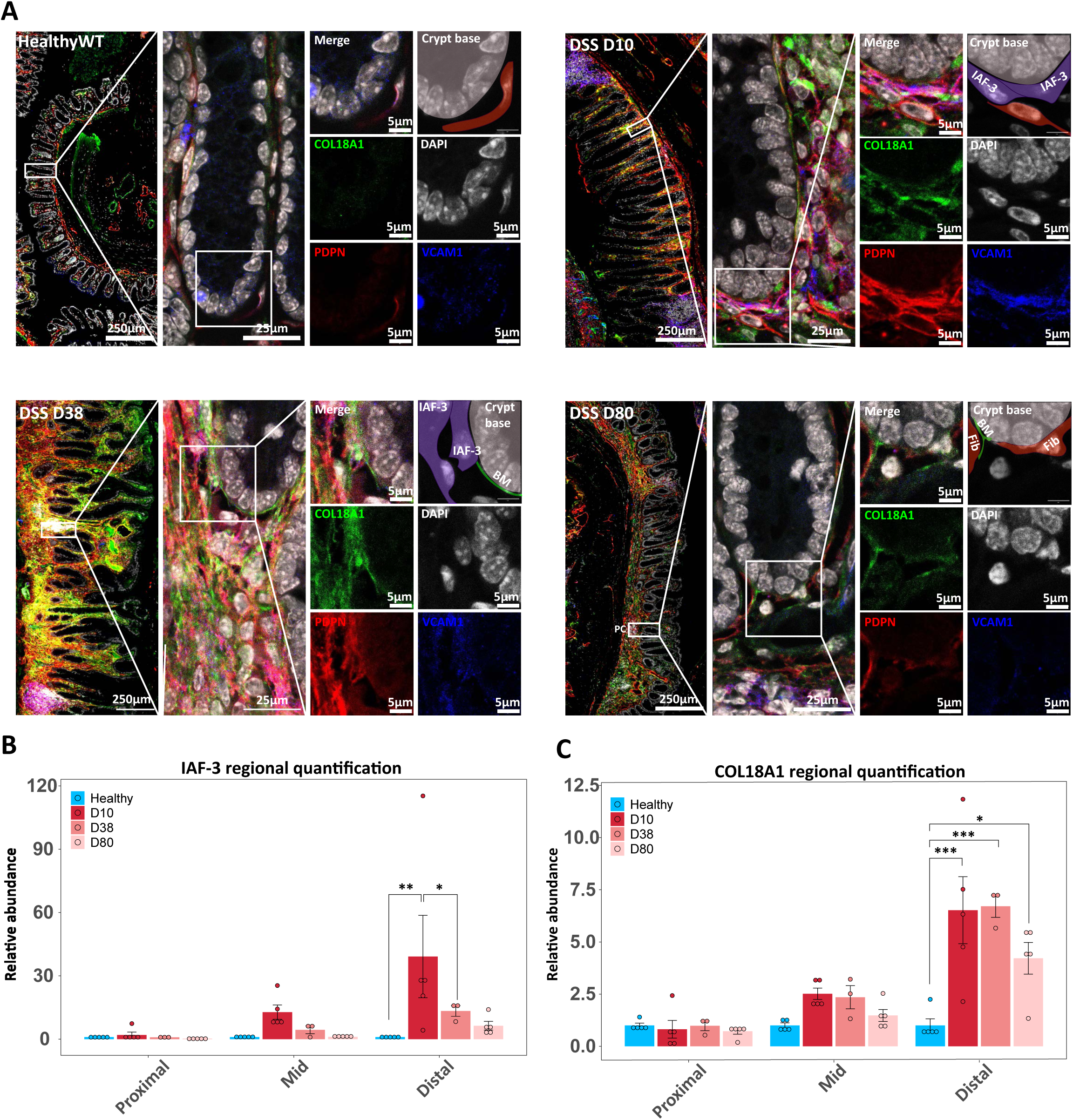
Inflammatory fibroblasts deposit COL18A1 in mucosal ECM. (**A**) Immunofluorescent images of pericryptal fibroblasts in the different DSS timepoints. Nuclei-White (DAPI), COL18A1-Green, PDPN-Red, and VCAM1-Blue. Each timepoint layout includes an image of repetitive crypts in the distal colon (left, scale bar of 250μm), magnified (mid, scale bar of 25μm), and split into its individual channels (right, scale bar of 5μm). Triple-positive fibroblasts are considered inflammation-associated fibroblasts (IAFs). The top right of each layout includes a simplified scheme of the different components in the image. BM, base membrane; Fib, fibroblast, IAF-3, Inflammation-associated fibroblast, subset 3^7^ (**B,C**) Relative quantification of IAF-3+ (B) and COL18A+ (C) covered area along the colon. *\* p≤0.05, ** p≤0.01, *** p≤0.001, log transformed two-way ANOVA, n ≥*3 mice per group.

These findings underscore the need for a deeper characterization of the ECM to uncover its dynamic changes. To this end, we initiated longitudinal mapping of decellularized ECM (dECM) composition to identify key signaling proteins and ECM-associated factors and their remodeling patterns over time.

### Longitudinal analysis of colon decellularized ECM (dECM) in the inflammatory and recovery phases

To further examine the ECM composition during inflammation and recovery phases, we extracted colon tissues from DSS-treated (Days 10, 38, and 80) and control (healthy WT) and decellularized the mucosal ECM (dECM) with minimal disruption of its structure (see **Methods**, **Figures 5A** and **S4A-C**), followed by LC-MS/MS proteomics. Using this technique, we enriched matrisome components while effectively removing non-tethered cellular and circulating proteins from the ECM during extraction. Of the identified 5710 differentially expressed proteins, 264 were matrisome proteins, with 64 proteins changing on Day 10, 23 proteins on Day 38, and 75 on Day 80 post-DSS treatment, compared to healthy controls **(Figure 5B, Table S3)**. Principal component analysis reveals that the composition of matrisome proteins in the dECMs distinctly differentiates between the healthy, acute, and regenerative phases **(Figure 5C)**. The identified altered matrisome proteins include core structural glycoproteins associated with matrix remodeling proteins. Consequently, as the matrix forms the foundation of every tissue, it also embeds numerous signaling molecules, growth factors, and small molecules such as cytokines and chemokines, serving as a reservoir for tissue signaling^28^. Interestingly, we identified numerous matrisome-bound proteins that exhibited significant changes during the inflammatory and recovery phases. Many of these proteins are associated with signaling pathways that regulate cell proliferation, cell fate, and response to environmental cues^28,45,46^. The mucosal dECM extracted during the regeneration period contained lower levels of core interstitial and basement membrane collagens, such as COL1A1, COL1A2, COL5A1, COL4A1, COL4A3, and higher levels of the squamous basement membrane COL7A1 and the pro-inflammatory proteoglycan COL18A1 **(Figure 5D-E)**. Laminin glycoproteins also constitute the basal membrane. The Lama1 subunit, a Laminin-111 component mostly present during embryogenesis and early epithelial development^47,48^, was increased in the regenerating colons. Conversely, the subunits (Lama3, Lamb3, and Lamc2) of Laminin-332 are key components for epithelium homeostasis by sustaining basement membrane assembly, cell adhesion, polarity, proliferation, and differentiation^49^, which were decreased **(Figure 5F)**. The significant loss of fibrillar collagens, coupled with the persistent accumulation of the glycoprotein COL18A1 beyond the recovery phase, could lead to biomechanical alterations in the dECM following DSS-induced tissue injury and throughout the regeneration phases. To validate the biomechanical impact of the modified ECM, we used atomic force microscopy to assess the elastic modulus of the dECM in the injury and recovery stages (**Figure S4D**). Our results revealed a progressive change in dECM stiffness, culminating in a marked reduction by Day 80 (**Figure S4E**). This suggests that the degradation of fibrillar and basal membrane collagens, along with their aberrant recovery—evidenced by the accumulation of COL18A1—leaves the mucosal ECM in a persistently altered biomechanical state.

**Figure 5.**
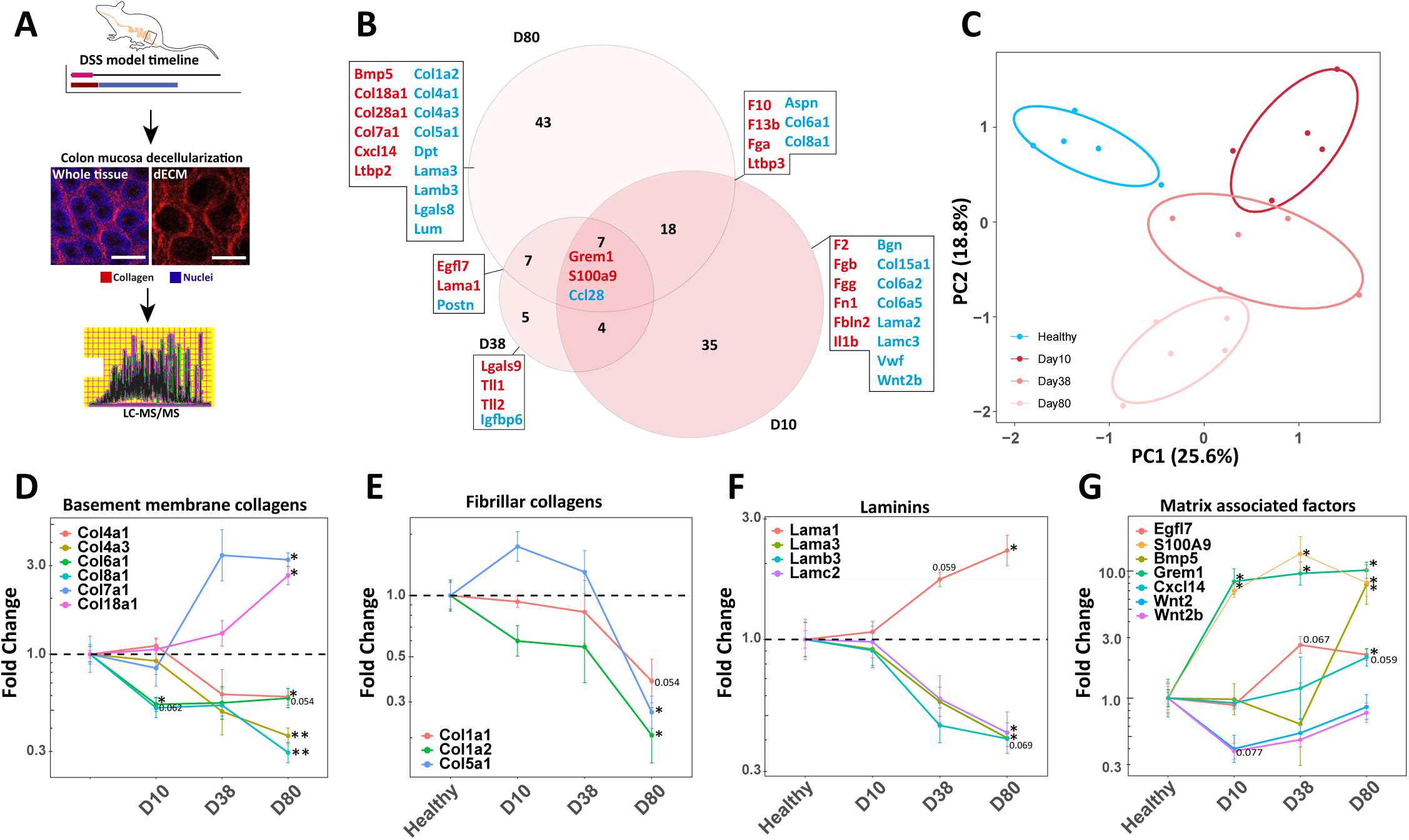
decellularized ECM composition during the DSS timepoints. (**A**) Scheme of colon extraction and decellularization followed by proteomics. (**B**) Venn diagram of matrisome proteins enriched in the different DSS-treated timepoints compared to healthy WT analyzed by LC-MS/MS (FDR < 0.1). Upregulated proteins are in red and downregulated in blue, compared to healthy colon (**C**) Principal component analysis (PCA) of decellularized matrisome from the DSS-induced colitis at different time points. Legends on the bottom left. (**D-G**) Trends of relative quantification (RQ) of matrisome proteins by type in the DSS timepoints, showing the gradual change from the acute Day 10 (D10) up to the restorative Day 80 (D80). Each color represents a different metrisome protein in the type. *\* p≤0.05, ** p≤0.01, FDR of log transformed student t test. n =* 5 mice per group.

In addition, we identified upregulated trapped signaling molecules in the dECM across the regenerative phases, such as the secreted pro-inflammatory factor S100a9, while the fibroblast chemokine, CXCL14, was elevated only in the last recovery phase **(Figure 5G)**. Other gut-related signaling molecules, such as EGFL7, BMP5, and its inhibitor GREM1, were detected at significantly higher levels. In contrast, WNT2 and WNT2b showed a non-significant decreasing trend **(Figure 5G)**. These results suggest that not only is the ECM structural composition modified, but also its protein composition, including key stem cell niche signaling molecules.

To further validate that the observed changes in the ECM-associated protein reservoir upon inflammation are not unique to the DSS-induced colitis model, we also examined the matrisome of dECMs extracted from Interleukin-10 knockout (IL-10 KO) mice, which experience spontaneous colitis around 5 months after birth^50^. We compared IL-10 KO mice with or without clinical symptoms to healthy WT mice of the same background (C57BL/6) and found that they too have distinct matrisomal compositions **(Figures S5A,B**), and the alterations in the matrisome from the healthy WT follow the same trend as observed in the DSS model **(Figure S5C, Table S3)**.

Overall, the inflammation-driven structural, compositional, and biomechanical modifications of the ECM, collectively termed “modECM,” create enduring damage imprints that persist beyond tissue recovery. More than mere remnants of past injury, these combinatorial alterations may actively shape cellular behavior and may serve as hidden instructors of tissue fate.

### modECM leads to intestinal stem cell pathogenic reprogramming

The profound and cumulative signals we found embedded within the modECM, reflected mostly in its altered composition and structure, may critically influence physiological epithelial restitution after injury. To test this hypothesis, we developed a two-component *ex vivo* model to investigate how the modECM might affect the colon epithelium, and in particular, the intestinal stem cells (ISCs), which play an important role in maintaining intestinal homeostasis via promoting a healthy gut barrier. We extracted dECMs from colon tissues under three conditions: healthy (control), acute inflammation phase (Day 10 post-DSS administration), and recovery phase (Day 38 post-DSS administration). dECMs were then suspended and embedded at the bottom of wells using GFR Matrigel as an adhesive (**Methods** and **Figure 6A**). Using spheroids conditioned media^51^ (**Methods**), cells grown on healthy dECM-enriched Matrigel formed spheroid-like colonies, whereas cells grown on dECM from Day 10 and 38 of DSS treatment were enriched with monolayers, throughout the entire 7-day experiment (**Figures 6B, C** and **S6A,C**). By Day 5, cells on inflamed dECM began spreading and losing their dome morphologies to form monolayers, while those on healthy dECM either retained their spheroid shapes or began budding into early organoids. By Day 7, fully developed organoids with complex structures emerged exclusively in cultures grown on healthy dECM, whereas cells on inflamed or recovered dECM remained in monolayer formations (**Figures 6B,C** and **S6A,C**). To confirm modECM is involved in the epithelial flattening observed in the DSS-associated dECMs, we utilized dECM preparations from the IL-10 KO model **(Figures S5D-F)**. In agreement with the DSS model, ISC-derived epithelial cells grown on WT dECM-enriched Matrigel developed into organoid morphologies, whereas ISCs grown on chronically inflamed IL-10 KO dECMs primarily formed monolayers (**Figure S5F**). These results further indicate that modECM alters ISC fate *ex vivo*, potentially influencing their ability to undergo normal self-renewal and differentiation, hence disrupting intestinal homeostasis.

**Figure 6.**
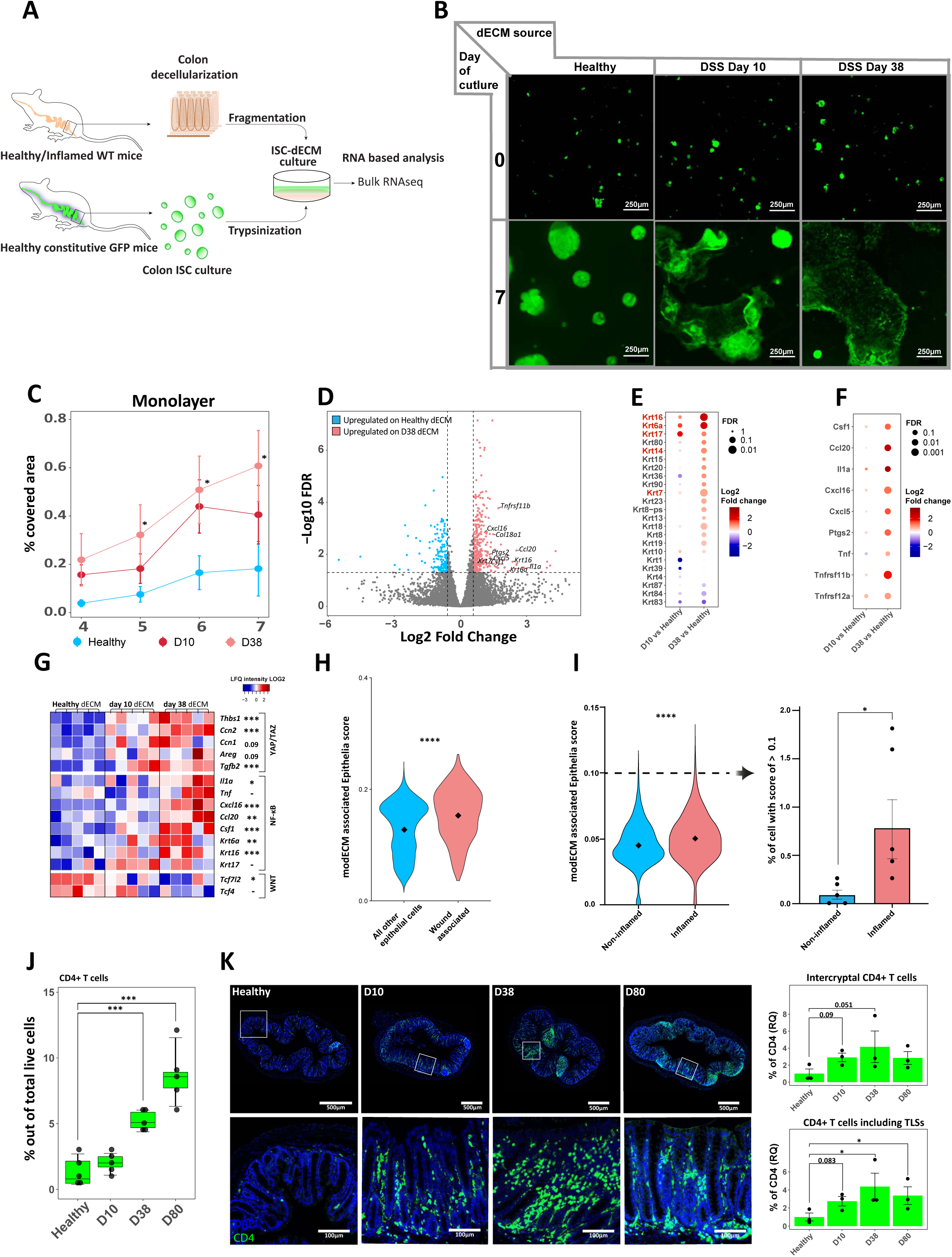
Characterization of modECM-associated epithelia transcriptome. (**A**) Scheme of the construction of the ISC-dECM culture assay. (**B**) Representative images of ISC-dECM cultures at Day 0 and 7 from different DSS-treated timepoints. Scale bar, 250μm. (**C**) Quantification of ISC-dECM monolayers. X axis, days after seeding. Y axis, ratio of covered area. Color of lines, healthy (blue), Day 10 (dark red), Day 38 (light red) dECM. *\* p≤0.1, one way ANOVA* comparing samples from the same time of culture, *n =* 5 mice (dECM samples) per group. (**D**) Differentially expressed genes (DEG) between ISCs grown on modECM (dECM from Day 38) compared to healthy dECM. Genes marked in red are significantly upregulated, and in blue downregulated on Day 38. (**E**) Cytokeratins are ordered by fold change of expression in ISCs grown on modECM Day 38 compared to healthy dECM. Red cytokeratins are known to be associated with wound-associated epithelia (WAE). The size of the dot indicates FDR significance, and the color indicates fold change expression. (**F**) Dotplot of proinflammatory cytokines and chemokines expressed by modECM-associated epithelia (DEG). The size of the dot indicates FDR significance, and the color indicates fold change expression. (**G**) Heatmap of key genes from signal transduction pathways identified in the GSEA pathway analysis. (**H**) Expression of dECM-associated epithelia signature in WAE cells (red) or all other epithelial cells (blue) from a previously published single-cell dataset of wound-associated epithelium^31^ (see **Methods**). *\*\*\*\* p≤0.0001, student t. test. n =* 5 mice per group. (**I**) Expression of dECM-associated epithelia signature in inflamed (red) and non-inflamed (blue) rectal biopsies of UC patients obtained from previously published single-cell data^27^ (see **Methods**). *\*\*\*\* p≤0.0001, student t. test. n =* 5 paired biopsies per group. (**J**) CD4+ T cell abundance in the colon was measured by cyTOF analysis within the DSS timepoints. *\*\*\* p≤0.001, log transformed one way ANOVA, n = 5* mice per group. (**K**) Representative IF staining of CD4+ T cells (green) and DAPI (blue), scale bar, 500μm. Inset, scale bar 100μm. Relative quantification (RQ) of intercryptal CD4+ cells (top right) and in all tissue (bottom right) *\* p≤0.05, log transformed one way ANOVA. n = 3 mice per timepoint*.

To gain mechanistic insights into the observed morphological changes, we performed bulk RNA sequencing of epithelial cells (**Table S4**) to identify genes associated with epithelial morphological changes induced by modECM compared to normal ECM. Our results show that ISCs grown on modECM (Day 38) enriched Matrigel induced pathophysiological cellular signaling compared to ISCs grown on the healthy dECM enriched Matrigel. To ensure there was no RNA carried over in the dECM samples, we compared the RNA integrity number (RIN) score of dECM samples alone to those with ISCs cultured and found that the RNA in dECM samples was completely degraded (**Figure S7A**). The RNA-seq analysis showed a significant increase in expression of pro-inflammatory cytokines and chemokines and squamous epithelial markers such as *Krt6a*, *Krt14, Krt16* in the modECM-epithelial cultures **(Figures 6D-F** and **Table S4)**. To better understand the gene programs associated with epithelial cells cultured on modECM (Day 38), we performed two pathway enrichment analyses—STRING enrichment and gene set enrichment analysis (GSEA) (**Figures S7B,C**). STRING identified enrichment in immune recruitment pathways (e.g., “T-cell migration”) and a decrease in the epithelial-cell commitment pathway. The GSEA identified the reduction in the “*Wnt* beta-catenin signaling”, crucial to maintain normal ISCs homeostasis ^52^ and inflammatory transcriptional pathways (e.g., “*Tnfα* signaling via *Nfκb*”) **(Figures S7B,C).** Indeed, the RNA-seq data revealed changes in three major transcriptional pathways– the canonical *Wnt* signaling, the *NF-κB*, and the Hippo (*Yap/Taz*) pathways (**Figure 6G**). Comparing Day 38 to healthy samples, the canonical Wnt transcription factors, *Tcf7l2* and *Tcf4* were downregulated, while the Hippo pathway target genes, *Thbs1, Ctcg, Cyr61, Areg,* and *Tgfβ2* ^53^ and *NF-κB* pathway target genes, such as *Tnf, Ccl20, Csf1*^54^ were upregulated. These changes were previously reported to occur during epithelial restitution ^55^. Notably, *in vivo* mouse models (DSS and IL-10KO) have been reported to promote the conversion of columnar epithelial cells to a squamous-like program associated with a wound-healing phenotype ^36^. To assess the similarity between our modECM-associated epithelial cells to WAE, we compared WAE transcriptional program identified by single cell analysis^31^ to the modECM-epithelial cells (**Methods** and **Figure 6H**). Strikingly, our dECM-associated epithelial cell signature was highly enriched in the examined WAE cells, suggesting that dECM-associated epithelial cells closely resemble WAE cells not only in morphology but also in their transcriptional program. The association of IBD with the appearance of WAE was shown before^56^. Squamous-like cells have been reported in severe colitis patients ^57–59^, but the role of modECM in converting ISC has not been previously demonstrated or discussed. Thus, due to our *ex vivo* findings, we asked whether ISC conversion into wound-associated epithelia-like cells (WAE) occurs in ulcerative colitis (UC) patients. To investigate this, we analyzed a previously published single-cell dataset ^27^ containing colon epithelial cell subsets from UC patients, including samples from both inflamed and non-inflamed tissues. Using the *ex vivo* modECM-associated epithelia signature, which includes WAE markers (e.g., *Ptgs2, Cxcl16, Aldh1a3, Krt16, Krt6a*), we identified a statistically significant enrichment of the modECM-associated epithelia signature in inflamed UC tissue (**Methods** and **Figure 6I**). Notably, this signature was detected exclusively in inflamed rectal biopsies, indicating that epithelial conversion to modECM epithelia is more prevalent in inflamed rectal tissue, mirroring our observations in the mouse model. Selection of cells with high score of dECM-associated epithelia in rectal samples of UC patients revealed around 8-fold increase in the number of these cells in inflamed compared to non-inflamed tissue (**Figure 6I**).

In addition, our pathway enrichment analyses further revealed that ISCs exposed to recovered dECM activated inflammatory signaling pathways associated not only with wound healing but also with immune cell recruitment. Notably, genes involved in T cell chemotaxis and activation were upregulated (**Figures 6D,F**), suggesting that the persistent abnormal modECM composition in the recovered tissue sustains a pro-inflammatory microenvironment. To validate the T cell migration into the tissue of modECM, we measured CD4+ T helper cells, known to be involved in the progression of IBD^60^, using CyTOF, FACS, and IFA, and showed massive accumulation of CD4+ T cells long after the DSS treatment (**Figures 6J,K**). We next characterize the different CD4+ T cell subsets within the tissue, showing a significant increase in T regulatory cells (Tregs), both thymic and peripheral, and TH17 associated with IBD progression and a significant reduction in TH1, associated with bacterial infection^38,61,62^ (**Figure S7D,E**). These findings demonstrate that even after inflammation subsides, the modECM retains the capacity to drive pathological epithelial remodeling and subsequently immune activation, potentially contributing to long-term tissue dysfunction and disease recurrence.

### Spatially resolved modECM delineates pathological regenerated colon

COL18A1, a key ECM component, has been identified as an early pre-symptomatic marker of IBD^20^, but its role in the post-injury regeneration and participation in cellular crosstalk remained elusive. Given our findings that modECM contains COL18A1-enrich regions in the distal colon (**Figure 4C**), we sought to investigate the modECM cellular crosstalk that shapes the pathological regenerative landscape of inflammation. Specifically, we performed spatial co-localization of COL18A1, the pan-immune cell marker CD45, and the modECM-epithelial cells marker KRT14 to define the pathological regenerative zone of the colon **(Figure 7A)**. Inspection of the different regions of the colon (proximal, mid, and distal) using colon Swiss rolls, showed the highest enrichment of COL18A1, CD45, and KRT14 positive signal in the distal colon and the transition zone (TZ) of the rectum (**Figures 7B-D)**. Of note, COL18A1 was not detected within TLSs, even due enriched in the DSS-treated colons, and suggests that non-immune cells are the main source of COL18A1 expression. Our data support that the major source of COL18A1 is inflammatory-associated fibroblasts (IAF) or modECM epithelial cells (**Figures 4** and **6D**, starting at Day 10). In addition, we observed sporadic KRT14^+^ cells emerging in the cryptal epithelium of the distal colon (around 5% at the late time points, **Figures 7E,F)**, in the proximity of COL18A1-rich regions in the pathological regenerative or recovery phases. These observations, along with our *ex vivo* cultures, led us to hypothesize that modECM niche signals drive the conversion of columnar Lgr5+ ISC into modECM-associated epithelia, characterized by squamous-like morphology. These cells may emerge as continuous stratified squamous cells in anorectal regions or as sporadic squamous-like WAE cells in the distal colon (**Figure 7F**), aligning with the enrichment of modECM-associated epithelia features observed in UC patients (**Figure 6I**). To test this hypothesis, we performed a fate mapping experiment using the ISC-specific Lgr5-CreER^T2^ Rosa26:TdTomato transgenic mice^63^ **(Figure 7G)**. After tamoxifen injections, TdTomato expression was observed in all colonic crypts and their progeny, whereas the squamous epithelium at the anal transitional zone remained silent in WT mice **(Figure 7H)**. We induced colitis using DSS and compared healthy control mice to DSS-treated mice on Day 38. We found patches of stratified squamous cells exhibited a strong TdTomato signal, indicating their origin from colonic crypt-derived Lgr5+ progeny **(Figures 7H-J)**. Immunofluorescent staining confirmed that these cells were KRT14^+^, thus indicating that colonic ISCs or their progeny were converted to TZ squamous-like cells and contributed to the squamous region’s expansion **(Figure 7J)**. These results suggest that modECM leads to pathogenic ISC conversion and subsequently to immune infiltration as measured by the enrichment of the triple-positive COL18A1, KRT14, and CD45 in the regenerated colon tissue, perpetuating gut inflammation.

**Figure 7.**
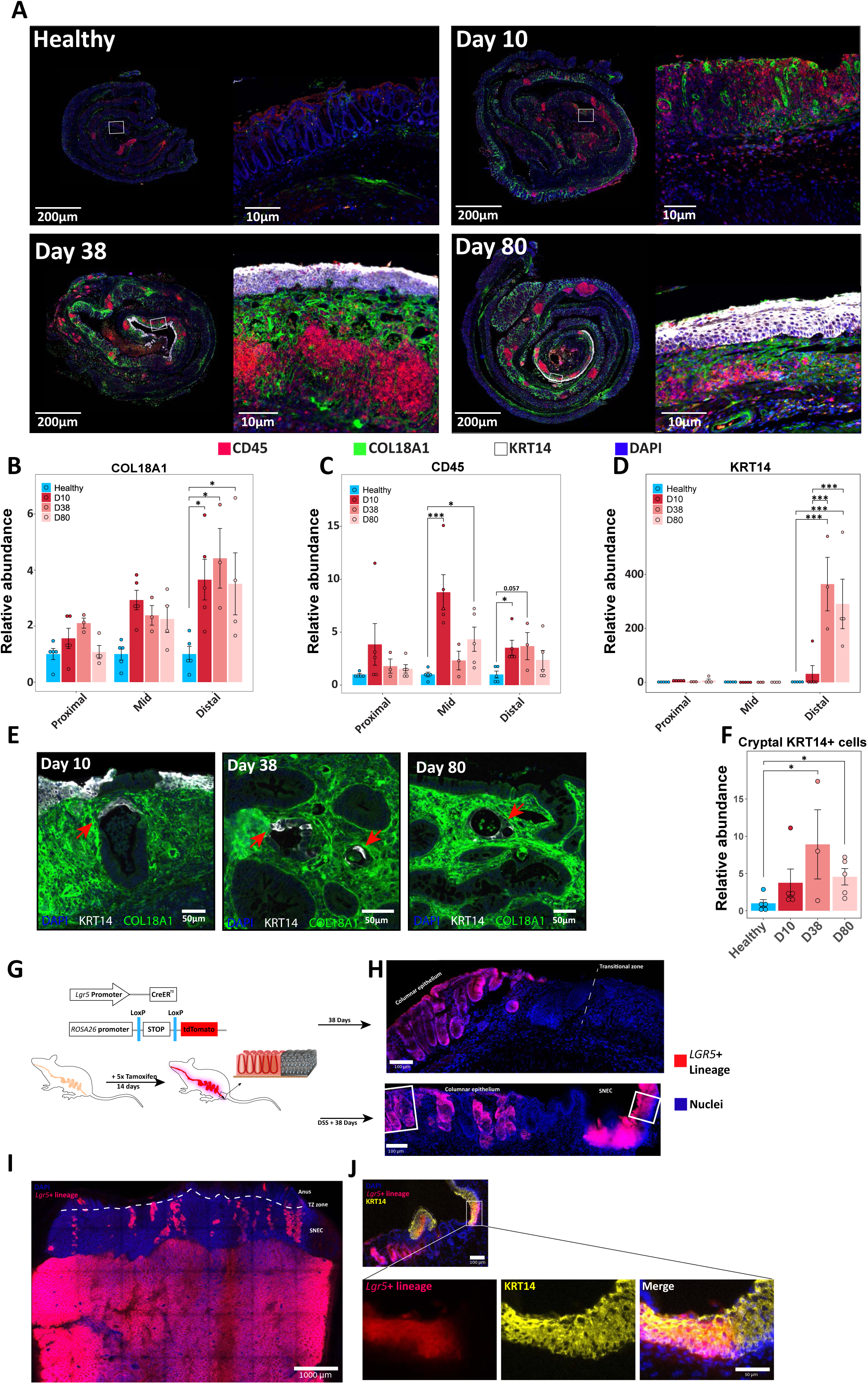
Identification of dECM regions and conversion of Lgr5+ ISC to modECM-associated epithelia. (**A**) Immunofluorescent staining of CD45+ (red) and KRT14+ (white) cells and COL18A1 (Green) of a representative whole colon from the DSS treatment timepoints. Scale bar, 200μm, Inset, 10μm. (**B-D**) Relative quantification of COL18A1 (B), CD45 (C), KRT14 (D) in different colon regions by percentage of covered area compared to healthy colon (WT). *\* p≤0.05, *** p≤0.001, log transformed two-way ANOVA, n ≥ 3* mice per group. (**E**) Representative IFA image of KRT14+ cells (white) emerged from colonic crypts in COL18A1-rich sites (green) in the distal colon from the DSS timepoints. Scale bar 50μm. (**F**) Relative quantification of modECM-associated epithelia percentage in colon mucosa (KRT14+ cells). (**G**) Scheme of fate mapping experiment. Lgr5-creERT2 crossed to Rosa26-lsl-tdTomato mice were used to follow Lgr5+ ISC fate. (**H**) Representative image of distal colon at anorectal transitional zone, healthy (top) and Day 38 of DSS treatment (bottom). Pink, Lgr5+ ISC progeny cells, DAPI for nuclei stain, scale bar 100μm. (**I**) Whole mount of distal colon giving an overview of Lgr5+ ISC progeny (pink) patches in SNEC. scale bar 1000μm. (**J**) Immunofluorescent staining of colocalization of KRT14+ (yellow) and Lgr5+ ISC progeny (Pink), scale bar, 100μm. Insets below, showing higher magnification of the co-localized cells, scale bar 50 μm.

## Discussion

Our study highlights a critical gap in the current understanding of extracellular matrix (ECM) remodeling and its crosstalk with stem cells during the regenerative and recovery phases following tissue damage. Traditionally, research on ECM repair has predominantly focused on acute tissue damage, operating under the assumption that both the ECM and resident cells eventually revert to their pre-injury state. This predominant view suggests that ECM remodeling restores the tissue’s original structure, composition, and functionality, and subsequently restores normal homeostasis. However, the findings presented here challenge this assumption by demonstrating persistent and profound alterations in the ECM, resulting in multifaceted damage to the ECM, which we have termed modECM. Specifically, our results using a mouse model of experimental colitis illustrate that acute inflammatory injury irreversibly modifies the ECM within the colon mucosa. Despite apparent epithelial recovery, these modifications persist, characterized by sustained basal membrane disruptions mediated by myeloid-derived proteolytic activity and abnormal accumulation of collagen XVIII to form a persistent scar deposited by inflammatory fibroblasts and WAE-like epithelial cells. Proteomic, biomechanical, and imaging analyses further revealed substantial and lasting alterations in ECM composition, indicative of persistent damage beyond the regeneration phase, which we characterized by loss of fibrillar collagen and alteration of signaling factors such as WNT and BMP ligands (**Figure 5**), all needed to maintain normal homeostasis of the stem-cell niche. Accordingly, our data demonstrate that healthy ISCs, when cultured on decellularized ECM derived from injury recovered tissue, adopt an inflammatory transcriptional profile and transition toward a wound-healing squamous-like epithelial phenotype, which we termed modECM-epithelia. Our results uncovered the importance of the colonic mucosal ECM niche to intestinal stem cells, particularly in the context of chronic diseases such as IBD, irrespective of genetic predisposition. Chronic conditions have long been characterized by cycles of remission and recurrence; however, the mechanisms that perpetuate disease recurrence remain mostly unknown. Our findings suggest that persistent ECM alterations, established during acute phases of inflammation, may serve as critical modulators of chronic disease progression by continuously influencing stem cell behavior long after clinical symptoms have subsided. Additionally, our study highlights the role of matrisome structural proteins and their embedded factors, which act as a reservoir of signals that directly impact pathological stem cell transitions.

Changes in the transcriptional expression of the ECM components, such as *Col18a1,* have been identified in a recent publication, peaking at the acute inflammation phase, and reduced with time, reflecting its cellular origin, the inflammatory-associated fibroblast subset 3 (IAF-3) abundance in the tissue^7^. We found that while IAF-3 cells are cleared during the recovery phase, their deposited COL18A1 is sequestered by the damaged ECM, contributing to the observed long-lasting modECM (**Figure 4**). Consequently, this transition, combined with ECM compositional, biomechanical and morphological modifications, alters the epithelial program and increases the presence of lymphoid lineages, particularly CD4+ T cells, which are associated with IBD (**Figure 6**). Specifically, we revealed that modECM pushes epithelial cells into pathogenic mode, characterized by pro-inflammatory transcriptional expression, with upregulation of cytokines and chemokines, such as *Cxcl16,* and *Cxcl5,* associated with T-cell recruitment.

Noteworthy, COL18A1 is a multifaceted heparin sulfate proteoglycan that does not form supramolecular fibrillar collagen structures ^64^. The COL18A1 domains are known to alter different signal transduction pathways with the potential to sequester Wnt ligands, disrupting Wnt gradient in the crypt-villus axis ^65^. The COL18A1 Heparan sulfate moiety could activate the NFκB^66,67^ and increase the ECM’s potential to bind a wide variety of chemokines, cytokines, and morphogens ^68–74^. Indeed, enriching cultures with dECM obtained from DSS recovered phases after acute injury induced the expression of the wound healing squamous-like epithelial program, we term modECM-associated epithelia, including upregulated YAP-signaling mediated fetal-like pathways (e.g., *Ly6a, Krt7, Anxa1, Krt14*) ^75,76^ and downregulated canonical Wnt pathway (e.g., *Tcf7l2, Tcf4, Fzd5*) (**Figure 6**). This modECM-associated epithelia program has previously been demonstrated in the context of wound healing, where wound-associated epithelium (WAE) emerged to facilitate the re-epithelialization of the colon^7,31,36,55^. In addition, recent studies showed the emergence of Krt14+ squamous neo-epithelial cells (SNECs), a form of WAE, in the distal colon ^36^. Our results imply that modECM, without any other cellular components, has the ability to trigger ISC conversion into KRT14+ WAE-like cells. Using COL18A1 as a marker for the modECM, we showed that KRT14+ cells can emerge from colon crypts engulfed with COL18A1 in the recovery phase (**Figure 7**) and found enrichment of the modECM-associated epithelial signature in inflamed rectal biopsies by querying UC patient single-cell data ^27^ (**Figure 6I**), suggesting the conversion of ISCs to modECM-associated epithelia under damage in the human context as well. Lastly, we utilized fate mapping of *Lgr5*-driven TdTomato to show that *Lgr5*^+^ ISCs contributed to modECM-associated epithelia *in vivo* via the conversion of ISCs into KRT14+ squamous-like epithelia in response to inflammatory cues.

Our findings emphasize an urgent and unmet need to broaden the scope of scientific inquiry regarding ECM-stem cell interactions during both regenerative and recovery phases of tissue healing. Importantly, further detailed investigations of ECM-stem cell dynamics throughout tissue regeneration and beyond can yield transformative insights, novel therapeutic targets, and improved patient outcomes in chronic disease management.

## Supporting information

Table S1 - DSS2 whole tissue proteomics only segnificant D38 vs Healthy

Table S2 - whole tissue degradomics only segnificant proteins comparing D38 to Healthy

Table S3 - dECM proteomics only segnificant genes comparing to healthy

Table S4 - dECM ISC culture RNAseq only segnificant genes comparing to healthy

Table S5 - Pathway analysis of RNAseq D38 vs Healthy

## Acknowledgement

We thank members of the Sagi laboratory for their advice and support of this work. We thank the Weizmann Core Facilities for their help and support, including Marina Cohen for histological work, Beni Siani, Oron Goresh and Shalom Faiz for animal care. Dr. Yoseph Addadi from the imaging unit. Muriel Chemla and Avital Sarusi-Portuguez for the genomic unit. Dr. Yishai Levin and Hila Levy from the Protein Profiling unit.

M.B. holds the Ernst and Kaethe Ascher Career Development Chair. This study was supported by research grants from the Center for New Scientists at the Weizmann Institute of Science, the Israel Science Foundation (grant No. 1587/20), the Helen and Martin Kimmel Institute for Stem Cell Research at The Weizmann Institute of Science, and the Minerva Foundation, with funding from the Federal German Ministry for Education and Research, the Moross Integrated Cancer Center (MICC), the Israel Ministry of Science (IMOS, Grant No. 4631), the Dr. Gilbert S. Omenn and Martha A. Darling Weizmann Institute - Schneider Hospital Fund for Clinical Breakthroughs through Scientific Collaborations, a research grant from the Snider Foundation, the Abisch-Frenkel RNA Therapeutics Center, the Shimon and Golde Picker grant, the Weizmann SABRA - Yeda- Sela - WRC Program, the Estate of Emile Mimran, and The Maurice and Vivienne Wohl Endowment Fellows, the Herbert K. Bennett Charitable Fund and Dwek Institute for Cancer Therapy Research. I.S. is an Incumbent of the Maurizio Pontecorvo Professorial Chair. I.S. research was supported by the Cynthia and Andrew Adelson Fund, the Rose Family Fund for Crohn’s and Colitis Research, the Israel Science Foundation (grant No. 1800/19), the Mireille & Murray Steinberg Family Foundation, the Thompson Family Foundation, and the Leonard and Carol Berall Foundation.

## Author contributions

M.B., I.S., and I.A. conceived the study, designed experiments, and interpreted the results; I.A. designed and performed the computational analysis with the assistance of B.T.; I.A., C.R., A.H.M., S.L., In.S. and, N.D. assisted in performing experiments; M.B. and I.S. supervised this study; T.M.S. assisted with CyTOF experiments and analysis; I.G. performed the atomic force microscopy analysis; S.S., F.S., and U.A.dK. assisted with proteomics and degradomic analysis; and I.A., I.S., and M.B. wrote the manuscript, with input from all authors.

## Supplemental Figures

**Figure S1 – DSS-induced colitis model characterization.**

(A) Inflammation score table assessed by colonoscopy^35^. (B) Representative colonoscopy images from the DSS-induced timepoints with their associated score. (C,D) Quantification of colon widths post-extraction with a representative image. *\* p≤0.05, log transformed one way ANOVA. n =* 4 mice per group. (E) Masson’s trichrome histochemical stain of DSS timepoints up to Day 400 with quantification of TLSs. *\* p≤0.05, ** p≤0.01, log transformed one way ANOVA, n ≥* 3 mice per group.

**Figure S2 – Cell type identification via cyTOF.**

Correlation heatmap displaying average intensity of the markers used for the cyTOF experiment per each population created in the tSNE graph shown in Figure 2A. EpCam (CD326) for epithelial cells, CD34 and CD31 for endothelial cells/progenitors (EEPs and LEPs), CD45 for immune cells, CD3E for T cells, where CD4 indicates T helpers and CD8 indicates cytotoxic T cells. CD45R (B220) is a marker of B cells, CD11C and MHC class II for dendritic cells, CD11b for myeloid cells, LY6G for neutrophils, LY6C (F4/80 Neg) for Monocytes, F4/80 for macrophages where iNOS+ for pro-inflammatory macrophages and CD64+ for the rest. Values are presented as scaled mean intensity values, see legend on top right.

**Figure S3 – Inflammation-associated fibroblasts in inflammation sites.**

Nuclei-White, COL18A1-Green, PDPN- Red, and VCAM1- Blue. Triple-positive fibroblasts are considered inflammation-associated fibroblasts (IAFs)^7^. The bottom right of each layout includes a simplified scheme of the different components in the image. (A) Image of ulceration, which is only present in the acute DSS Day 10 timepoint. (B,C) Images of the squamous and the expanded lamina propria below crypts in the recovered DSS Day 38 and 80 timepoints. Scale bars are depicted on the images.

**Figure S4 – ECM decellularization and alteration during colitis progression.**

(A) Representative images of colon mucosa throughout the decellularization process. (B,C) IF and SHG imaging of Z projection of whole tissue mount (B) and decellularized ECM (dECM) sample (C) demonstrating the removal of cells in the decellularization process. Red-fibrillar collagen (SHG), Green-EPCAM, Blue-PDPN, White-Nuclei. Huygens-SVI deconvolution v.24.10 was used to remove noise. (D) Representative images of intercryptal dECM measurements using atomic force microscopy (AFM). (E) Stiffness measurement of dECM via atomic force microscopy. Units according to young’s modulus (kPa). *\*\* p≤0.01, log transformed student’s t-test n≥*5 mice per group.

**Figure S5 – IL-10 Knockout induced colitis.**

(A) Scheme of the IL-10 KO mouse model as used in this experiment. (B) Principal component analysis (PCA) of decellularized matrisome from the IL-10 KO mouse model. Legens on the top left. (C) Matrisome comparison between the fold change of the inflamed IL-10 KO vs Healthy WT (y-axis) and DSS Day 80 compared to WT (x-axis). The correlation between IL-10 KO and DSS 80 Day is depicted on the graph (r2=0.386). Legends is on the bottom right. (D) Representative image of ISC-dECM culture on day 7, using the dECMs extracted from IL-10 KO mouse model (non-inflamed vs inflamed). GFP spheroids were collected from healthy GFP transgenic mice. Scale bar, 500 μm. (E, F) Quantification of ISC-dECM morphologies - small spheroids, large spheroids (area > 350μm), and budding organoids (E) or monolayers (F). X axis, days after seeding. Y axis, ratio of counts of the chosen morphologies out of all identified bodies. Color of lines, healthy (blue), IL-10 KO inflamed (dark red), IL-10 KO non-inflamed (light red) dECM. *\* p≤0.05, ** p≤0.01, *** p≤0.001, one way ANOVA* comparing samples from the same time of culture, *n =* 5 mice (dECM samples) per group.

**Figure S6 - Morphological image analysis and quantification ISC-dECM culture.**

(A,B) Each row in each table contains replicates of either healthy ISCs cultured on dECM fragments from Healthy WT or DSS treatment (Day 10 and Day 38) (A) or IL-10 KO (non-inflamed or inflamed) (B). All images are after 7 days of culture (n ≥ 5 biological replicates per group). The image analysis pipeline classifies the morphologies in these images into: small spheroids (circularity ≥ 0.61-1.0, area ≤ 350μm), large spheroids (circularity ≥ 0.85, area > 350μm), organoids (circularity ≥ 0.7, size > 350μm) and monolayers (circularity < 0.7 area ≤ 700μm). Images were taken using the EVOS M500 with 4X/0.13 lens. Scale bar, 1mm. (C) Quantification of ISC-dECM morphologies - small spheroids, large spheroids (area > 350μm), and budding organoids. X axis, days after seeding. Y axis, ratio of counts of the chosen morphologies out of all identified bodies. Color of lines, healthy (blue), Day 10 (dark red), Day 38 (light red) dECM. *\* p≤0.05, ** p≤0.01, *** p≤0.001, one way ANOVA* comparing samples from the same time of culture, *n =* 5 mice (dECM samples) per group.

**Figure S7 – dECM-associated epithelia transcriptome and CD4+ T cell subsets *in vivo*.**

(A) Rin score of dECM samples with ISCs (first 4 from the left marked in green) or without ISCs (marked in orange). Rin score below 3 indicates RNA is of very low integrity. (B) Gene enrichment analysis using STRING with Gene Ontology gene sets (left), or GSEA (gene set enrichment analysis). Red marks pathways upregulated in ISCs grown on modECM Day 38, and in blue are pathways upregulated in ISCs grown on healthy WT dECM. (C) GSEA output graphs corresponding to enrichment scores in (B). (D-E) Cytometry analysis of DSS timepoints of CD4+ T cells extracted from colon. (D) Shows the gating strategy used to define each population. Antibodies used for gating are depicted on the bottom left of each column. (E) Relative quantification (RQ) of CD4+ T subsets out of all CD45+ immune cells. *\* p≤0.05, *** p≤0.001, log transformed one way ANOVA, n =* 5 mice per group.

**Figure S1 – DSS-induced colitis model characterization.**

(A) Inflammation score table assessed by colonoscopy^35^. (B) Representative colonoscopy images from the DSS-induced timepoints with their associated score. (C,D) Quantification of colon widths post-extraction with a representative image. *\* p≤0.05, log transformed one way ANOVA. n =* 4 mice per group. (E) Masson’s trichrome histochemical stain of DSS timepoints up to Day 400 with quantification of TLSs. *\* p≤0.05, ** p≤0.01, log transformed one way ANOVA, n ≥* 3 mice per group.

**Figure S2 – Cell type identification via cyTOF.**

Correlation heatmap displaying average intensity of the markers used for the cyTOF experiment per each population created in the tSNE graph shown in Figure 2A. EpCam (CD326) for epithelial cells, CD34 and CD31 for endothelial cells/progenitors (EEPs and LEPs), CD45 for immune cells, CD3E for T cells, where CD4 indicates T helpers and CD8 indicates cytotoxic T cells. CD45R (B220) is a marker of B cells, CD11C and MHC class II for dendritic cells, CD11b for myeloid cells, LY6G for neutrophils, LY6C (F4/80 Neg) for Monocytes, F4/80 for macrophages where iNOS+ for pro- inflammatory macrophages and CD64+ for the rest. Values are presented as scaled mean intensity values, see legend on top right.

**Figure S3 – Inflammation-associated fibroblasts in inflammation sites.**

Nuclei- White, COL18A1- Green, PDPN- Red, and VCAM1- Blue. Triple-positive fibroblasts are considered inflammation-associated fibroblasts (IAFs)^7^. The bottom right of each layout includes a simplified scheme of the different components in the image. (A) Image of ulceration, which is only present in the acute DSS Day 10 timepoint. (B,C) Images of the squamous and the expanded lamina propria below crypts in the recovered DSS Day 38 and 80 timepoints. Scale bars are depicted on the images.

**Figure S4 – ECM decellularization and alteration during colitis progression.**

(A) Representative images of colon mucosa throughout the decellularization process. (B,C) IF and SHG imaging of Z projection of whole tissue mount (B) and decellularized ECM (dECM) sample (C) demonstrating the removal of cells in the decellularization process. Red- fibrillar collagen (SHG), Green-EPCAM, Blue-PDPN, White-Nuclei. Huygens-SVI deconvolution v.24.10 was used to remove noise. (D) Representative images of intercryptal dECM measurements using atomic force microscopy (AFM). (E) Stiffness measurement of dECM via atomic force microscopy. Units according to young’s modulus (kPa). *\*\* p≤0.01, log transformed student’s t-test n≥*5 mice per group.

**Figure S5 – IL-10 Knockout induced colitis.**

(A) Scheme of the IL-10 KO mouse model as used in this experiment. (B) Principal component analysis (PCA) of decellularized matrisome from the IL-10 KO mouse model. Legens on the top left. (C) Matrisome comparison between the fold change of the inflamed IL-10 KO vs Healthy WT (y- axis) and DSS Day 80 compared to WT (x-axis). The correlation between IL-10 KO and DSS 80 Day is depicted on the graph (r2=0.386). Legends is on the bottom right. (D) Representative image of ISC-dECM culture on day 7, using the dECMs extracted from IL-10 KO mouse model (non-inflamed vs inflamed). GFP spheroids were collected from healthy GFP transgenic mice. Scale bar, 500 μm. (E, F) Quantification of ISC-dECM morphologies - small spheroids, large spheroids (area > 350μm), and budding organoids (E) or monolayers (F). X axis, days after seeding. Y axis, ratio of counts of the chosen morphologies out of all identified bodies. Color of lines, healthy (blue), IL-10 KO inflamed (dark red), IL-10 KO non-inflamed (light red) dECM. *\* p≤0.05, ** p≤0.01, *** p≤0.001, one way ANOVA* comparing samples from the same time of culture, *n =* 5 mice (dECM samples) per group.

**Figure S6 - Morphological image analysis and quantification ISC-dECM culture.**

(A,B) Each row in each table contains replicates of either healthy ISCs cultured on dECM fragments from Healthy WT or DSS treatment (Day 10 and Day 38) (A) or IL-10 KO (non-inflamed or inflamed) (B). All images are after 7 days of culture (n ≥ 5 biological replicates per group). The image analysis pipeline classifies the morphologies in these images into: small spheroids (circularity ≥ 0.61-1.0, area ≤ 350μm), large spheroids (circularity ≥ 0.85, area > 350μm), organoids (circularity ≥ 0.7, size > 350μm) and monolayers (circularity < 0.7 area ≤ 700μm). Images were taken using the EVOS M500 with 4X/0.13 lens. Scale bar, 1mm. (C) Quantification of ISC-dECM morphologies - small spheroids, large spheroids (area > 350μm), and budding organoids. X axis, days after seeding. Y axis, ratio of counts of the chosen morphologies out of all identified bodies. Color of lines, healthy (blue), Day 10 (dark red), Day 38 (light red) dECM. *\* p≤0.05, ** p≤0.01, *** p≤0.001, one way ANOVA* comparing samples from the same time of culture, *n =* 5 mice (dECM samples) per group.

**Figure S7 – dECM-associated epithelia transcriptome and CD4+ T cell subsets *in vivo*.**

(A) Rin score of dECM samples with ISCs (first 4 from the left marked in green) or without ISCs (marked in orange). Rin score below 3 indicates RNA is of very low integrity. (B) Gene enrichment analysis using STRING with Gene Ontology gene sets (left), or GSEA (gene set enrichment analysis). Red marks pathways upregulated in ISCs grown on modECM Day 38, and in blue are pathways upregulated in ISCs grown on healthy WT dECM. (C) GSEA output graphs corresponding to enrichment scores in (B). (D-E) Cytometry analysis of DSS timepoints of CD4+ T cells extracted from colon. (D) Shows the gating strategy used to define each population. Antibodies used for gating are depicted on the bottom left of each column. (E) Relative quantification (RQ) of CD4+ T subsets out of all CD45+ immune cells. *\* p≤0.05, *** p≤0.001, log transformed one way ANOVA, n =* 5 mice per group.

## Methods

### Animal models

All mouse work was performed in accordance with the Institutional Animal Care and Use Committees (IACUC, no. 05050623-3, 02130222-2) of the Weizmann Institute of Science. Mice were housed under specific-pathogen-free conditions at the animal facilities at the Weizmann Institute of Science, Rehovot, Israel. C57BL/6J wild type (WT) mice were purchased from Envigo. The DSS-induced colitis model is done on 8-week-old WT or transgenic C57BL/6 male mice. Drinking water with 1.25% DSS (MP Biomedicals LLC) was provided to the mice. After 7 days, their water was switched back to normal water ^33^. Colitis progression is evaluated throughout the experiment using the Karl Storz Coloview mini-endoscope system. IL-10 KO mice (B6.129P2-Il10tm1Cgn/J) and GFP- expressing mice (C57BL/6-Tg(UBC-GFP)30Scha/J) were purchased from Jackson Laboratories and inbred locally. The Lgr5-CreER^T2^, kindly provided by Prof. Nick Barker from the A*STAR Institute of Molecular and Cell Biology, Singapore, was crossed to the ROSA26-lsl-TdTomato mouse strain to generate a fate mapping of Lgr5+ ISCs. To induce Cre recombinase activation and lineage tracing, mice were injected 5 consecutive times with Tamoxifen (40 mg/Kg per injection).

### Mucosal separation and tissue decellularization

Whole colons were harvested, carefully separated from the surrounding fat and cleaned from feces with cold PBS. Then, the colons were cut with a longitudinal incision to expose the colon lumen. Colon samples were incubated for 1 hour in 10 mM EDTA PBS x1 on ice. After the incubation, loose epithelial cells were rinsed with PBS, and the muscle layer was peeled off from the mucosal layer with forceps under a surgical microscope.

Mucosal layers were then placed in new 15 mL tubes and incubated for 24 hours in the first decellularization buffer (50 mM buffer Tris (pH 8.0), 50 mM EDTA, 1.5 M NaCl with 1:50 tablet of cOmplete protease inhibitor, 1% Triton) at 4°C. Samples were then washed 3 times in DW and placed in the second decellularization buffer (1% deoxychalate in DW) for 1 hour at RT. Samples were then washed in DW 3 times and incubated with 0.1 mg/mL DNase in PBS with calcium chloride for 1 hour at 37°C. Samples were then incubated in penicillin/streptomycin 10x (BI 03-031-1B), supplemented with 0.5 mg/mL gentamicin (Sigma G1272-10ML) overnight at 4°C.

### dECM proteomics

Decellularized tissues (about 50mg per tissue) were submerged in hard tissue homogenization tubes in a solution of 10 mM EDTA, 10 mM DTT, 100 mM HEPES, and 1% SDS (1 mg/5uL concentration). The derived dECMs were then homogenized in a bead beater for 4 intervals of 30 seconds at 4°C. Lysates are then transferred to 1.5 mL tubes and sonicated on high for 10 minutes. Next, 1 µL of benzonase (Sigma) was added, followed by incubation on ice for 20 minutes. Tubes were centrifuged at 4°C at 10,000 RCF for 10 minutes. After the BCA protein concentration evaluation, 10 µg was taken for Methanol/Chloroform precipitation. Samples were then resuspended in 10 µL of 0.1 M NaOH, supplemented with 5 µL of HEPES pH 7.3 and 35 µL of GuHCL 6 M. Samples were reduced with TCEP 10 mM (final concentration) and alkylated with chloroacetamide 40 mM (final concentration). Next, 350 µL of HEPES pH 7.3 and trypsin 1:100 (trypsin(mg):protein(mg)) were added and incubated overnight at 37°C by preferably shaking. Samples were then acidified with 1 µL trifluoroacetic acid, stage-tipped, and dehydrated using a Speedvac. Samples were analyzed with LC-MS/MS on the HFX mass spectrometer (Thermo). The proteins were identified by the Proteome Discoverer 2.4 (Thermo) vs. the mouse UniProt database. The detected peptides are filtered for proteins identified with FDR < 0.05. Statistical analysis was finally performed using RStudio. Differentially expressed proteins were determined using *student T- test* on log2 protein values followed by *benjamini-hochberg* false discovery rate (FDR).

### Degradomics

For degradomics analysis of colon tissue from DSS time points, we performed terminal amine isotope labeling of substrates (TAILS) experiments, n ≥ 3 biological replicates per each time point, were snap frozen in liquid nitrogen. Prior to protein denaturation and labeling, proteins of the samples were extracted by mechanical homogenization using Tissuelyser (QIAGEN) and sonication using Bioruptor Pico (Diagenode). This process was done on 10 mg tissue in a buffer containing 4 M Guanidine hydrochloride (Sigma-Aldrich), 250 mM HEPES (Sigma-Aldrich (pH 7.8)) and protease inhibitor cocktail (cOomplete EDTA-free, Roche).

TAILS analysis, whereby protein N-termini were enriched, was performed according to the previously described protocol^77^, except for using 16 plex Tandem mass Tag™ labeling (TMT, #90,110, Thermo Fisher) and adding the trypsin at a ratio of 1:20 (trypsin/protein ratio).

PreTAILS and TAILS peptide mixtures were resuspended in MS injection buffer (1%TFA, 2%Acetonitrile, containing iRT peptides (Biognosys)) loaded onto an PepMap100 C18 precolumn (75 μm × 2 cm, 3 μm, 100 Å, nanoViper, Thermo Scientific) and separated on a PepMap RSLC C18 analytical column (75 μm × 50 cm, 2 μm, 100 Å, nanoViper, Thermo Scientific) using an EASY- nLC™ 1200 liquid chromatography system (Thermo Scientific) coupled in line with a Q Exactive HF-X mass spectrometer (Thermo Scientific). Separation of 500 ng of peptide mixture was achieved by running a constant flow rate of 250 nL/min in 0.1% formic acid/99.9% water and a 140 min gradient from 10% to 95% (23% for 85 min, 38% for 30 min, 60% for 10 min, 95% for 15 min) elution buffer (80% acetonitrile, 0.1% formic acid, 19.9% water). A data–dependent acquisition (DDA) method operated under Xcalibur 3.1.66.10 was applied to record the spectra. For the preTAILS and TAILS samples the full scan MS spectra (350–1500 m/z) were acquired with a resolution of 120′000 after accumulation to a target value of 3e6 in profile mode. MS/MS spectra with fixed first mass of 110 m/z were recorded in centroid mode from the 12 most intense signals with a resolution of 45,000 applying an automatic gain control of 1e5 and a maximum injection time of 96 ms. Precursors were isolated using a window of 0.7 m/z and fragmented with a normalized collision energy (NCE) of 30%. Precursor ions with unassigned, single charge or above charge of six were rejected, and precursor masses already selected for MS/MS were excluded for further selection for 60 s. Settings for the scan of the secretome peptide mixture were the same except for using a resolution of 17′500 for recording the MS/MS spectra, a maximum injection time of 60 ms, and a NCE of 25%.

Raw files were searched against the Mus musculus database compiled from the UniProt reference proteome (ID: 10,090, entries: 17′005) using Sequest from Proteome Discoverer™ 2.4 software (Thermo). The following parameters were selected for database searches: semi-ArgC for enzyme specificity with tolerance of one missed cleavage; carbamidomethyl(C) and TMTpro (K) as fixed modifications, and acetyl(N-term), pyroQ (N-term), TMTpro (N-term), oxidation(M), deamidation (NQ), as variable modifications. Precursor mass error tolerance of 10 ppm and fragment mass error at 0.02 Da. Percolator was used for decoy control and FDR estimation (0.01 high confidence peptides, 0.05 medium confidence). For protein–level analysis of the preTAILS sample, quantifiable peptides were summed up per corresponding protein.

Further statistical analysis (namely, hierarchical clustering and heatmap generation) was done on Log2 values of the protein intensities. Statistical analysis was performed using RStudio. Differentially expressed proteins were determined using *student T-test* on log2 protein values followed by *benjamini-hochberg* false discovery rate (FDR). The data is available at ProteomeXchange Consortium (http://www.proteomexchange.org/) via the PRIDE partner repository with the dataset identifier PXD022668.

### Atomic force microscopy

AFM imaging and mechanical measurements were conducted with a JPK Nanowizard III AFM microscope (Bruker Nano GmbH, Berlin, Germany) in QI and force spectroscopy modes. In this mode, force–distance curves are recorded at each pixel, these are used to acquire topographic images simultaneously with nano-mechanical data. Measurements were conducted with a 30 µm glass colloid glued on a qp-BioAC- probe (Nanosensors, Neuchâtel, Switzerland), spring constant ≈ 0.2 N/m. Images of 50x50 µm2 were captured in 60x60 pixel resolution to locate the crypts walls, Force curves form the center of each crypt wall were used to calculate the elastic modulus, by applying contact mechanics Hertzian model with Poisson ratio of 0.5 and a spherical tip with 15 µm radius. The force applied on each pixel was 2 nN and the approach speed was 40 µm/s. Image analysis was performed using Gwyddion^78^ and JPK-SPM data processing software. (version 6.1.86).

### Primary ISC culture

Whole colons were harvested, carefully separated from the surrounding fat, and cleaned from feces with cold PBS. They were then minced and washed in PBS on ice. When no more fat pieces remained, tissue fragments were incubated for 2 hours in 10 mM EDTA in PBS x1 at RT. Samples were then centrifuged at 1000 RPM, aspirated, and washed with PBS twice. After the last centrifugation, the remaining pellet was resuspended in 280 µL growth factor-reduced matrigel, supplemented with Jag1, and plated in 30 µL domes. After the matrigel polymerizes, domes were cultured in stem cell enrichment media (50% base - DMEM/F12 with HEPES (GIBCO 12634-010) + 10% (GluM, FBS, 1x penicillin/streptomycin, 10 µM Y27632 (TOCRIS 1254) and 10 µM SB431542 (TOCRIS 1614)). Subculturing was done every 5 days.

### ISC-dECM culture set up model

ECM samples were fragmented by mixing decellularized tissue samples with PBS at a 1 mg to 5 µL ratio within a hard tissue homogenization tube. Tubes were then shaken at 4°C for 30 seconds at level 4 for 4 cycles. At the end of this process, each dECM was plated at a 1:1 ratio with growth factor- reduced matrigel in a 96-well plate (40uL per well). Plates were then centrifuged at 800 g for 3 minutes to pull down and condense the dECM and matrigel while also getting rid of bubbles. Plates were placed at 37°C overnight. Wells were then washed with PBS and incubated with 100 µL of the respective growth media (as described in the sections above). Suspended cells (10^5^) of the cell type of choice were then plated on top of the culture with an additional 100 µL of media. The cells were allowed to sink toward the dECM fragments for optimal spreading.

### Mass cytometry (CyTOF)

Colons were harvested, carefully separated from the surrounding fat, and cleaned from feces with cold PBS. Then, 2-cm colon pieces were minced and immersed into RPMI-1640 medium (GIBCO, 21875034) containing 10% FBS, 2 mM HEPES, 0.05 mg DNase (Roche, 04716728001), and 0.5 mg/mL of collagenase VIII (Sigma-Aldrich, C2139), incubated for 40 min at 250 rpm shaking at 37°C, and finally transferred through a 70 mm cell strainer. Cell suspensions from each mouse were stained with Cell-ID Cisplatin 1.25 μM (Standard BioTools, 201064) for viability and fixed using Maxpar Fix I Buffer (Standard BioTools, 201065). Samples were then permeabilized using Maxpar Barcode Perm Buffer (Standard BioTools, 20166) and then barcoded using The Cell-IDTM 20-Plex Pd Barcoding Kit (Standard BioTools, 201060), allowing us to join samples for antigen staining. The antibodies used for staining were labeled with lanthanides. Before analyzing, the cell suspension was incubated with Cell-ID Interculator Iridium 250nM (Standard BioTools, 201192A) for 20 min. Cells were analyzed with a CyTOF2 mass cytometer. Results were normalized and barcoded using the Fluidigm CyTOF software. CyTOF results gating^79^ and further analysis were done with the FlowJo software (FlowJo, LLC).

### CyTOF antibody panel

**CyTOF antibody panel:**
| *Commercial antibodies* | **Metal** | **Product#** | **Manufacturer** |
| --- | --- | --- | --- |
| *Ly-6G (1A8)* | 141Pr | 3141008B | Fluidigm |
| *CD11c (N418)* | 142Nd | 3142003B | Fluidigm |
| *CD11b M1/70* | 143Nd | 3143015B | Fluidigm |
| *CD16/32-93* | 144Nd | 3144009B | Fluidigm |
| *CD4 (RM4-5)* | 145Nd | 3145002B | Fluidigm |
| *CD8a (53-6.7)* | 146Nd | 3146003B | Fluidigm |
| *CD140a (APA5)* | 148Nd | 3148018B | Fluidigm |
| *Ly-6C- (HK1.4)* | 150Nd | 3150010B | Fluidigm |
| *CD64 (X54-5/7.1)* | 151Eu | 3151012B | Fluidigm |
| *CD3e (145-2C11)* | 152Sm | 3152004B | Fluidigm |
| *Foxp3 (FJK-16s)* | 158Gd | 3158003A | Fluidigm |
| *F4/80 (BM8)* | 159Tb | 3159009B | Fluidigm |
| *CD45R/B220 (RA3-6B2)* | 160Gd | 3160012B | Fluidigm |
| *TER-119-162Dy* | 162Dy | 3162003B | Fluidigm |
| *BCL-6 (K112-91)* | 163Dy | 3163012B | Fluidigm |
| *CD31/PECAM-1 (390)* | 165Ho | 3165013B | Fluidigm |
| *CD326/EpCAM (G8.8)* | 166Er | 3166014B | Fluidigm |
| *Gata3 (TWAJ)* | 167Er | 3167007A | Fluidigm |
| *Anti-Ki-67 (B56)* | 168Er | 3168007B | Fluidigm |
| *IL-17A (TC11-18H10.1)* | 169Tm | 3169005B | Fluidigm |
| *NK1.1 (PK136)* | 170Er | 3170002B | Fluidigm |
| *CD86 (GL1)* | 172Yb | 3172016B | Fluidigm |
| *I-A/I-E (M5/114.15.2)* | 209Bi | 3209006B | Fluidigm |
| *Anti-Human CD45 (HI30)* | 89Y | 3089003B | Fluidigm |

| ***Homemade conjugations*** | **Metal** | **Product#** | **Manufacturer** |
| --- | --- | --- | --- |
| *MMP9* | 149Sm | ab38898 | Abcam |
| *LOX* | 175Lu | ab31238 | Abcam |
| *MMP7* | 148Nd | ab5706 | Abcam |
| *ADAM-17* | 174Yb | ab57484 | Abcam |
| *MMP14* | 156Gd | ab51074 | Abcam |
| *iNOS* | 147Sm | ab3523 | Abcam |

#### Flow cytometry analysis

Colons were harvested, carefully separated from the surrounding fat, and cleaned from feces with cold PBS. Then, 2-cm colon pieces were minced and immersed into 5mL RPMI-1640 medium (GIBCO, 21875034) containing 2% FBS, 0.05 mg, DNase (Roche, 04716728001), and 150uL Liberase (Sigma-Aldrich, 5401119001), incubated for 1 hour at 250 rpm shaking at 37°C, and finally transferred through a 70 mm cell strainer. Immune cells were isolated via Percoll gradient (Sigma- Aldrich, P4937) using a 80% and 40% gradient and centrifuging at 1000g for 20 minutes at room temperature. Cells were then washed, incubated with TruStain FcX™ PLUS (Biolegend, 156604), and then stained with fluorescently labeled antibodies. Samples were analyzed with the Beckman CytoFLEX S. Results gating and further analysis were done with the FlowJo software (FlowJo, LLC).

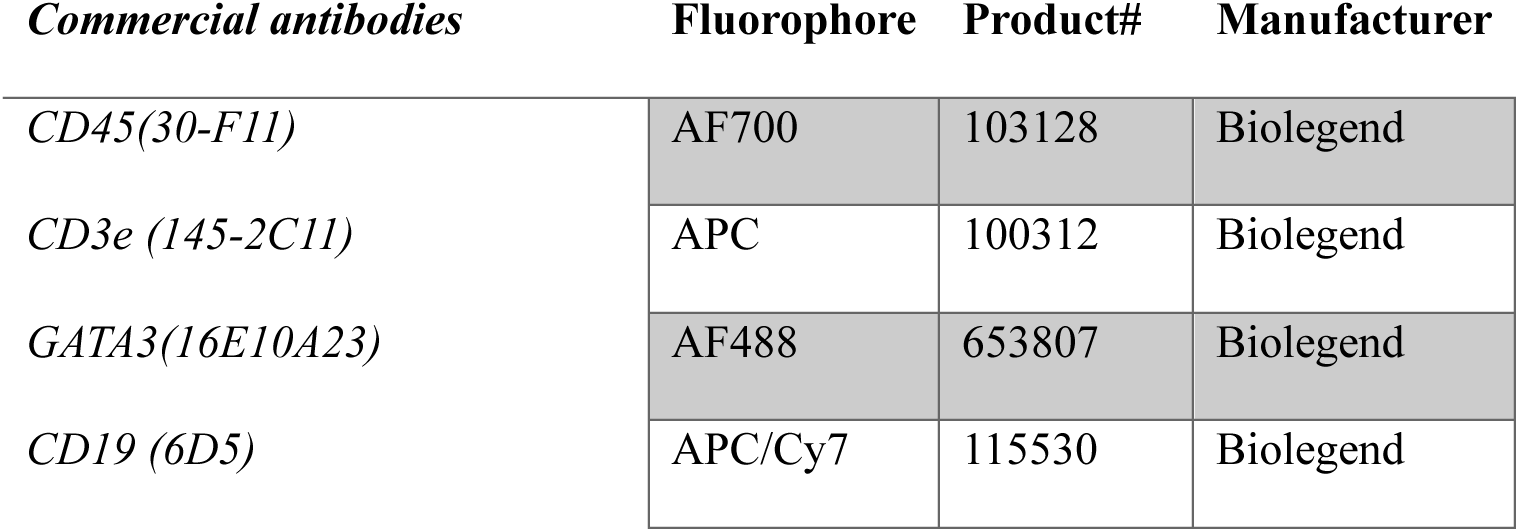
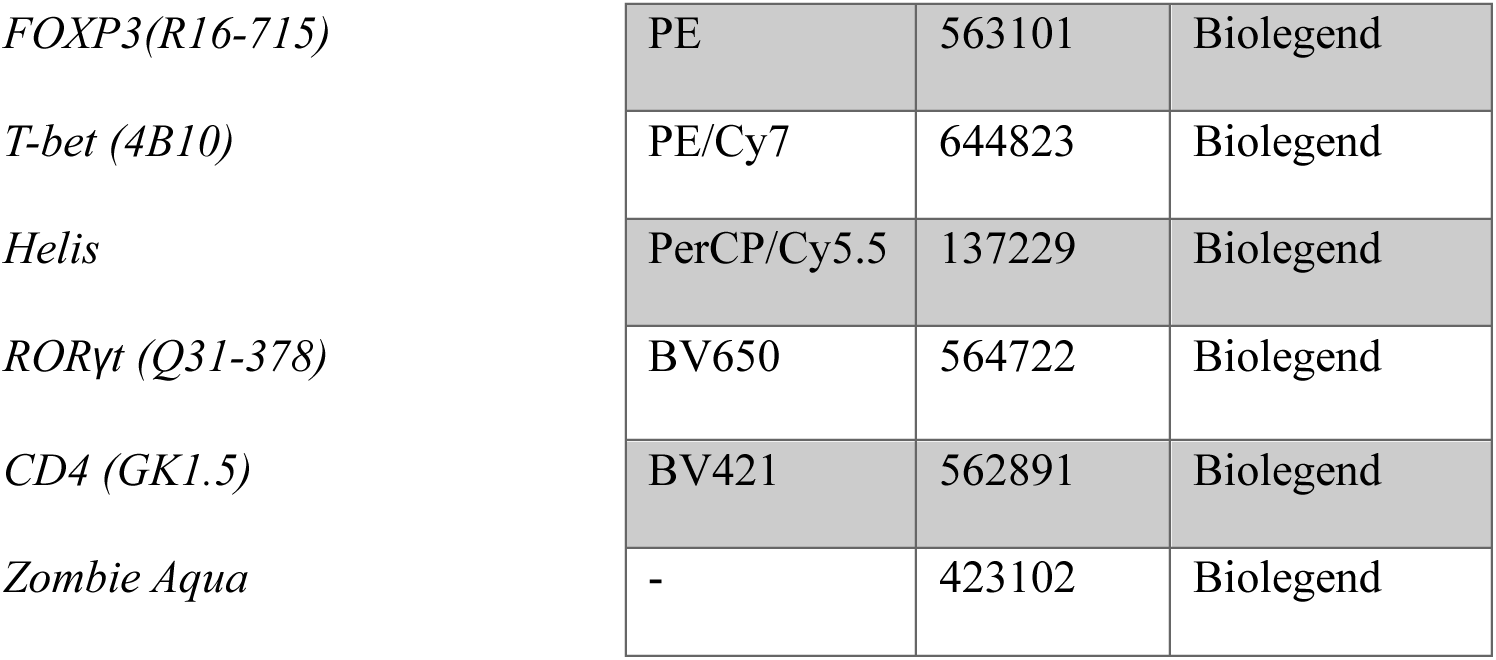

#### Immunofluorescence staining

Samples were fixed with 4% paraformaldehyde in PBS. The fixed samples were either (i) paraffin- embedded and sectioned (4 μM), or (ii) OCT-embedded, frozen on dry ice and stored at -80◦C, and then sectioned (50 μM). Paraffin Slides were de-paraffinized and subjected to antigen retrieval in Tris-EDTA or citric acid buffer. Samples were blocked in PBS, 20% normal donkey serum, and 0.1% Triton X-100 (60 min, 25◦C) and then incubated with primary antibodies in PBS, containing 2% normal donkey serum and 0.2% Triton X-100, overnight at 4◦C. Next, samples were washed three times in PBS and incubated with a secondary antibody (60 min, 25◦C), and mounted in a mounting medium. Image analysis and quantifications were done using ImageJ software.

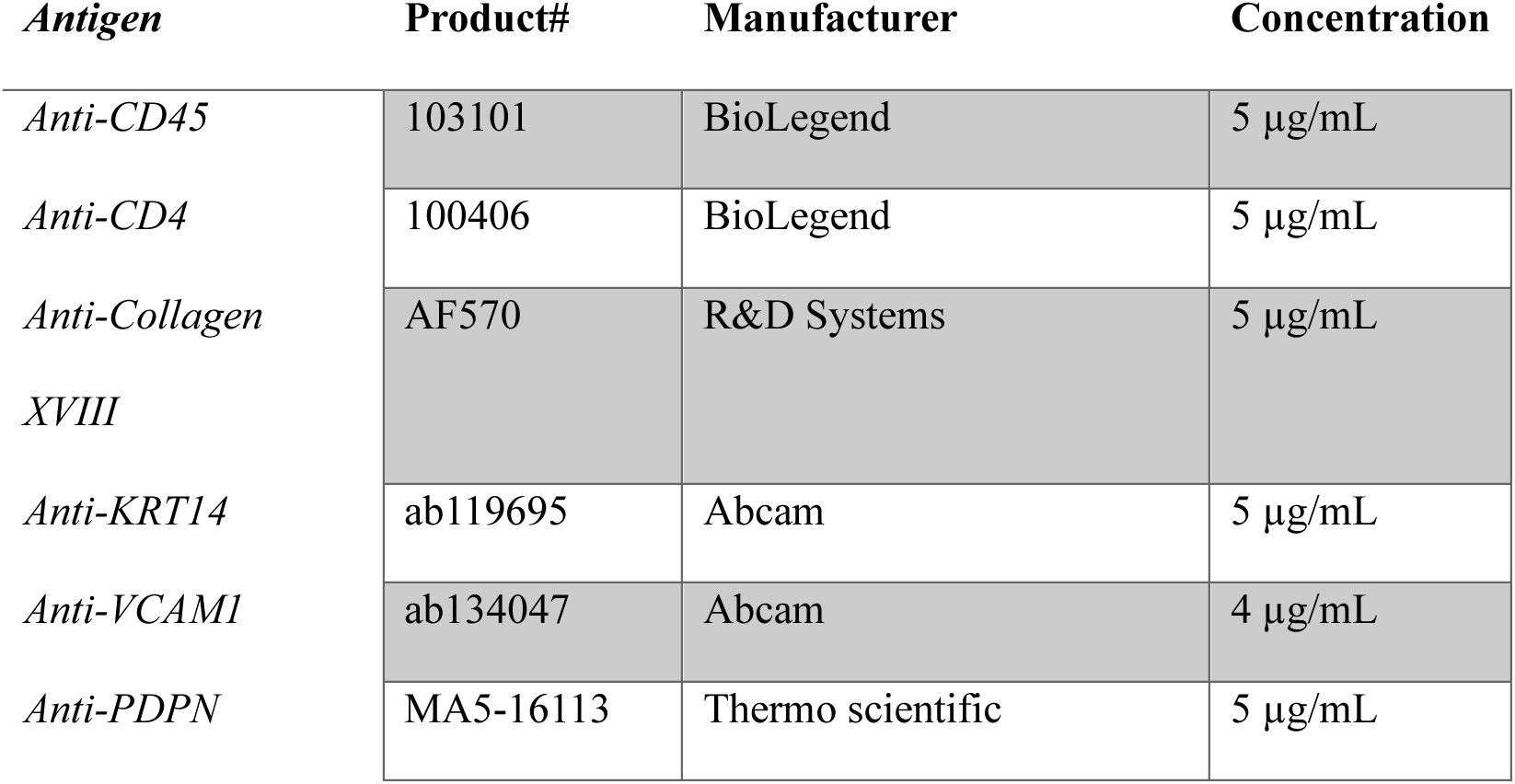

#### Second Harmonic Generation (SHG) module using two-photon microscopy

Native snap-frozen murine colon samples were thawed in PBS and imaged using a Leica TCS SP8 confocal multiphoton photon microscope. For second-harmonic imaging of collagen, a wavelength of 850 nm was used (detection at 390-450 nm). Image analysis is done using ImageJ and Amira for initial analysis and quantification.

#### RNA purification

Total RNA was isolated using the Tri-reagent/chloroform (sigma) separation technique according to the manufacturer’s protocol, and the aqueous top phase was taken for RNA purification.

#### Bulk RNA sequencing

Libraries were prepared using a modified SMART-Seq2 protocol^80^. RNA lysate cleanup was performed using RNAClean XP beads (Agencourt), followed by reverse transcription with Maxima Reverse Transcriptase (Life Technologies) and whole transcription amplification (WTA) with KAPA HotStart HIFI 2 3 ReadyMix (Kapa Biosystems) for 18 cycles. WTA products were purified with Ampure XP beads (Beckman Coulter), quantified with Qubit dsDNA HS Assay Kit (ThermoFisher). PE reads were sequenced on 1 lane(s) of an Illumina NovaSeq. The output was ∼11 million reads per sample.

Bioinformatics: Poly-A/T stretches, Illumina, and Nextera adapters were trimmed from the reads using cutadapt ^81^; resulting reads shorter than 25bp were discarded. Reads were mapped to the *M. musculus* reference genome GRCm39 using STAR ^82^, supplied with gene annotations downloaded from Ensembl (and with EndToEnd option and outFilterMismatchNoverLmax was set to 0.04). Expression levels for each gene were quantified using htseq-count ^83^, using the gtf above. Differentially expressed genes were identified using DESeq2 ^84^ with the betaPrior, cooksCutoff and independentFiltering parameters set to False. Raw P values were adjusted for multiple testing using the procedure of Benjamini and Hochberg. The pipeline was run using snakemake^85^.

Pathway analysis was done using (i) STRING ^86^ analysis coupled with the “protein with values/ranks” method and referring to results from the Gene Ontology database ^87,88^, and (ii) the Gene Set Enrichment Analysis (GSEA) ^89,90^ based on hallmark pathways for mice (MH).

#### Cell signature score analysis

To quantify modECM-associated epithelia signature within our previously published human single- cell RNA sequencing (scRNA-seq) dataset ^27^, or within a previously published dataset of wound- associated epithelia (WAE)^31^, we employed the UCell R package. The analysis was conducted according to the standard workflow described in the UCell documentation. The UCell package and its documentation are publicly available on GitHub at https://github.com/carmonalab/UCell ^91^.

#### Statistics and graphs

All analyses were performed using R Statistical Software (v4.5.0; R Core Team 2025, R Foundation for Statistical Computing, Vienna, Austria). Statistics that were not related to LC-MS/MS proteomics or RNA sequencing were done with one-way ANOVA if there was one categorical variable or two- way ANOVA if there were two categorical variables using glht R package^92^. Statistics were done on log-transformed values, unless there was a 0 value, in which case the original values were used. Graphs were made with the R package ggplot2^93^.

## Declaration of generative AI and AI-assisted technologies in the writing process

During the preparation of this work, the authors used ChatGPT in order to improve writing quality. After using this tool, the authors reviewed and edited the content as needed and take full responsibility for the content of the publication.

