## Supplemental figures for "Persistent ECM Scarring Reprograms Intestinal Stem Cells to Drive Chronic Inflammation"

Figure S1.

**A**

| Category \ Score | 0 | 1 | 2 | 3 |
|---|---|---|---|---|
| **Feces** | Solid or absent | Compressible or little remains | Spreadable | Fragmented |
| **Blood** | None | Few spots and hematomas | Many spots and hematoma | Hemorrhaging |
| **Opacity** | Clear | Translucent | Opaque and pink | Opaque and white |
| **Granularity** | Smooth | Shallow bumps | Many bumps but still circular | Non-circular lumen |
| **Fibrin** | None | Few deposits | Many deposits | Fibrin cross lumen |

| **Score** | 0-4 | 5-7 | 8-11 | 12-15 |
|---|---|---|---|---|
| **Inflammation level** | Healthy | Mildly inflamed | Inflamed | Severly inflamed |

**B**

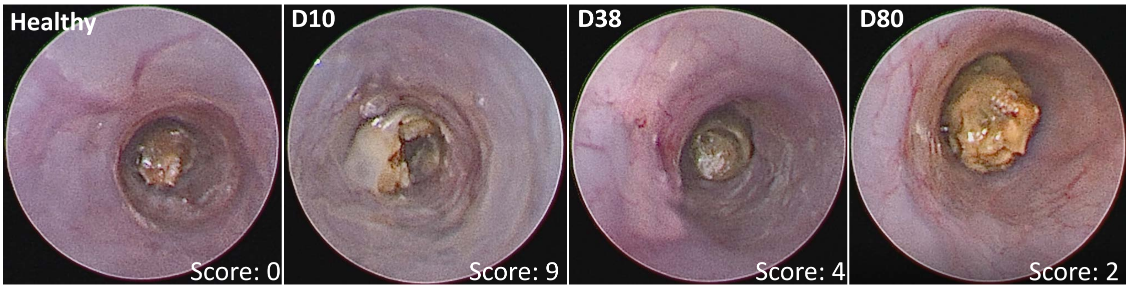


**C**

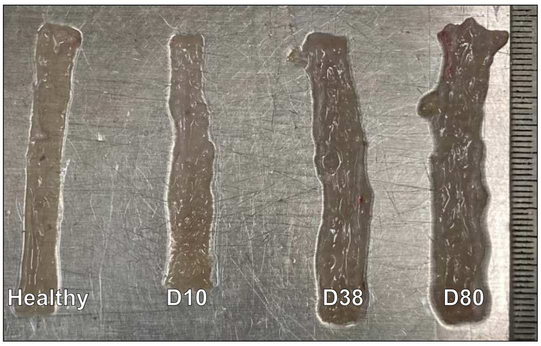


**D**

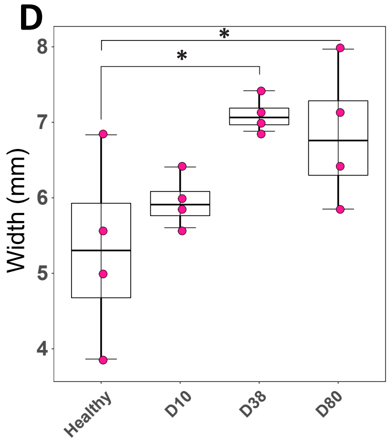


**E**

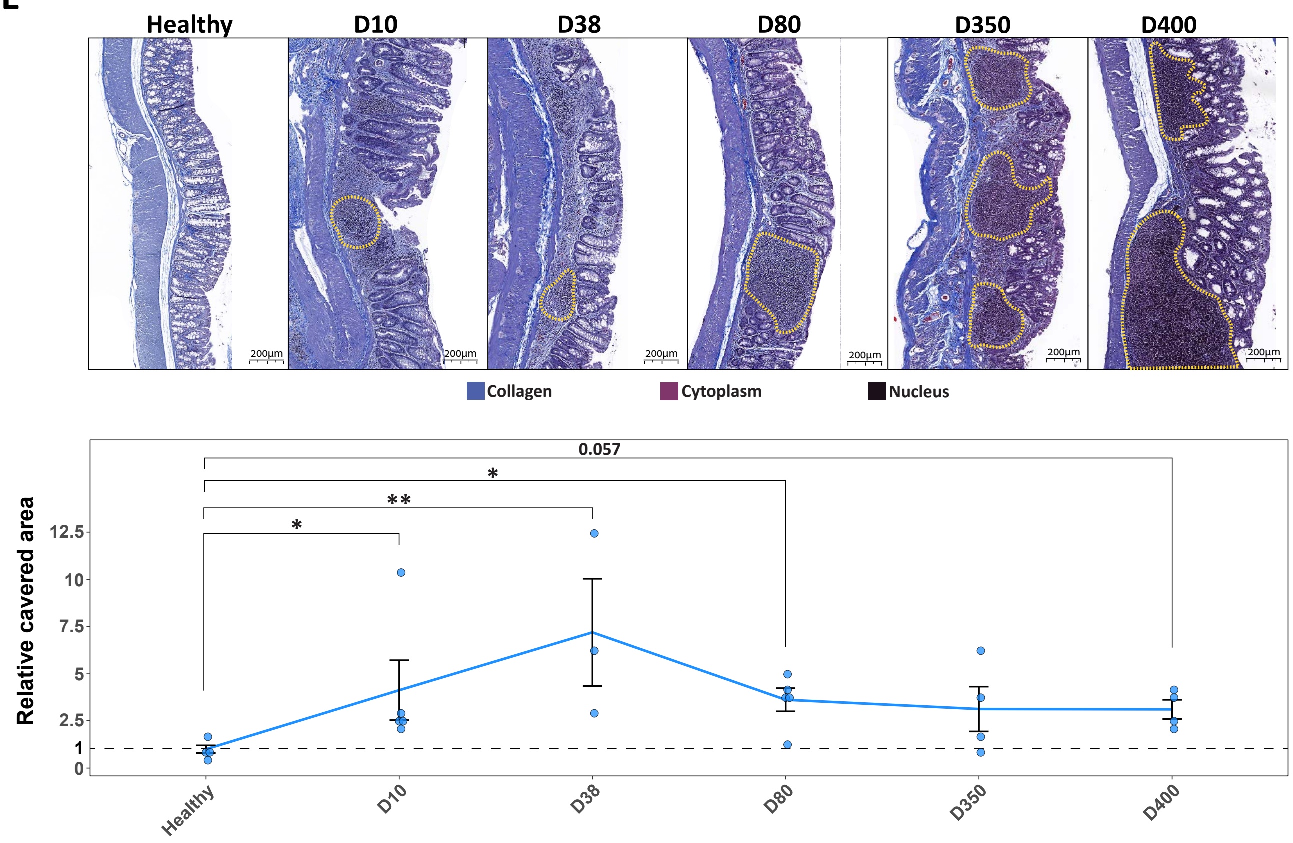

Figure S2.

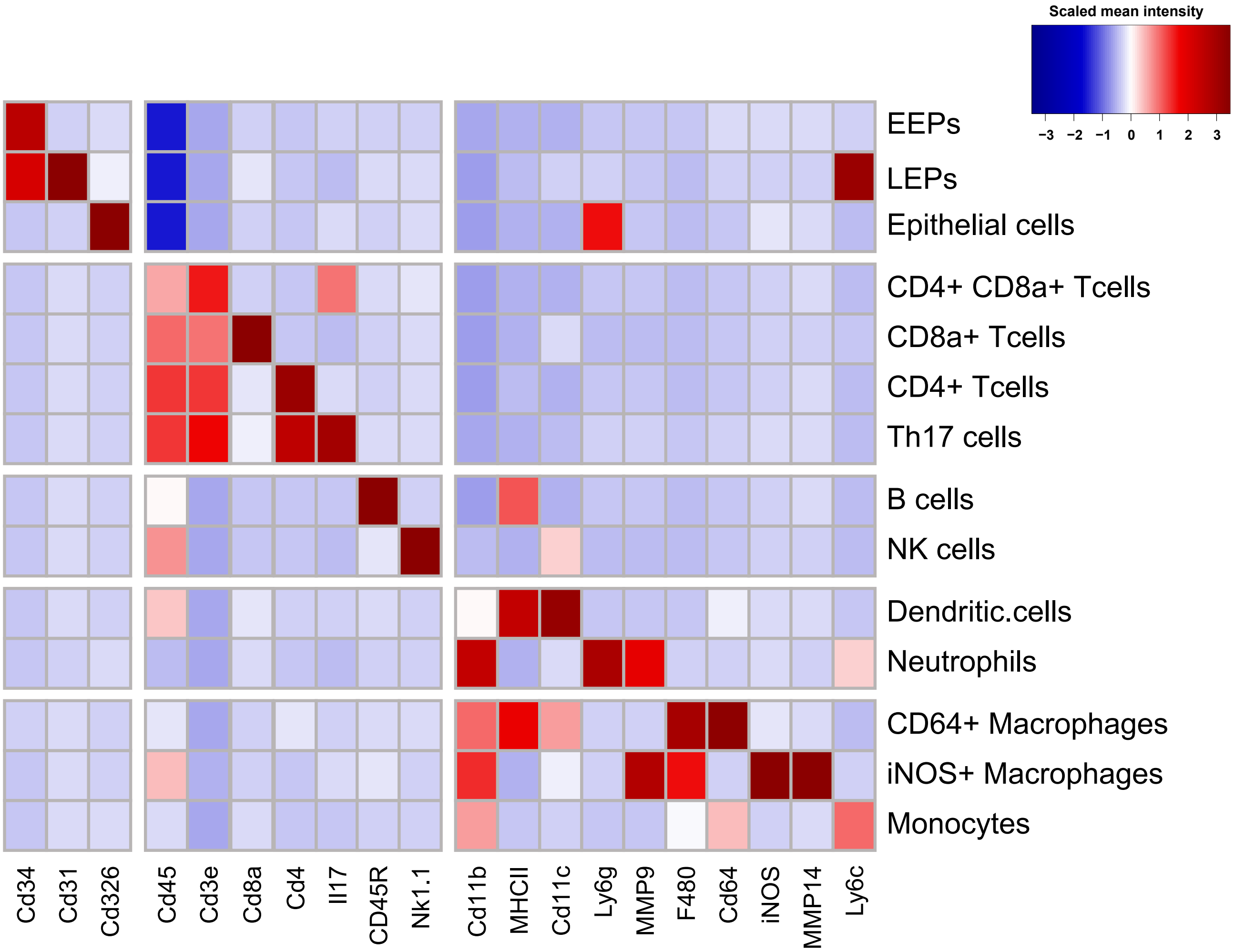

Scaled mean intensity
−3 −2 −1 0 1 2 3
EEPs
LEPs
Epithelial cells
CD4+ CD8a+ Tcells
CD8a+ Tcells
CD4+ Tcells
Th17 cells
B cells
NK cells
Dendritic.cells
Neutrophils
CD64+ Macrophages
iNOS+ Macrophages
Monocytes
Cd34
Cd31
Cd326
Cd45
Cd3e
Cd8a
Cd4
Il17
CD45R
Nk1.1
Cd11b
MHCII
Cd11c
Ly6g
MMP9
F480
Cd64
iNOS
MMP14
Ly6c

Figure S3.

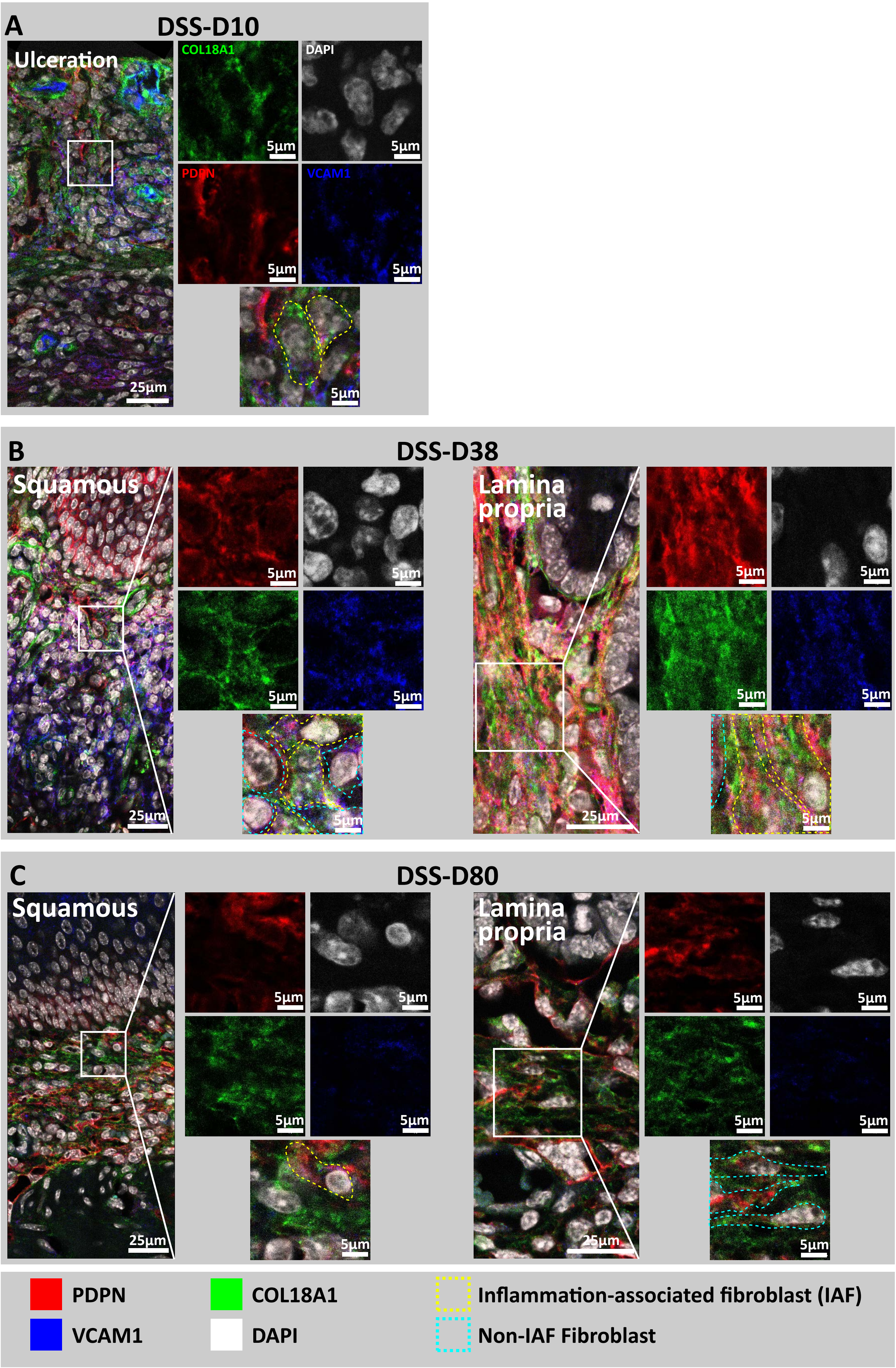
A
DSS-D10
Ulceration
COL18A1
DAPI
PDPN
VCAM1
5µm
25µm
B
DSS-D38
Squamous
Lamina propria
5µm
25µm
C
DSS-D80
Squamous
Lamina propria
5µm
25µm
PDPN
VCAM1
COL18A1
DAPI
Inflammation-associated fibroblast (IAF)
Non-IAF Fibroblast

Figure S4.

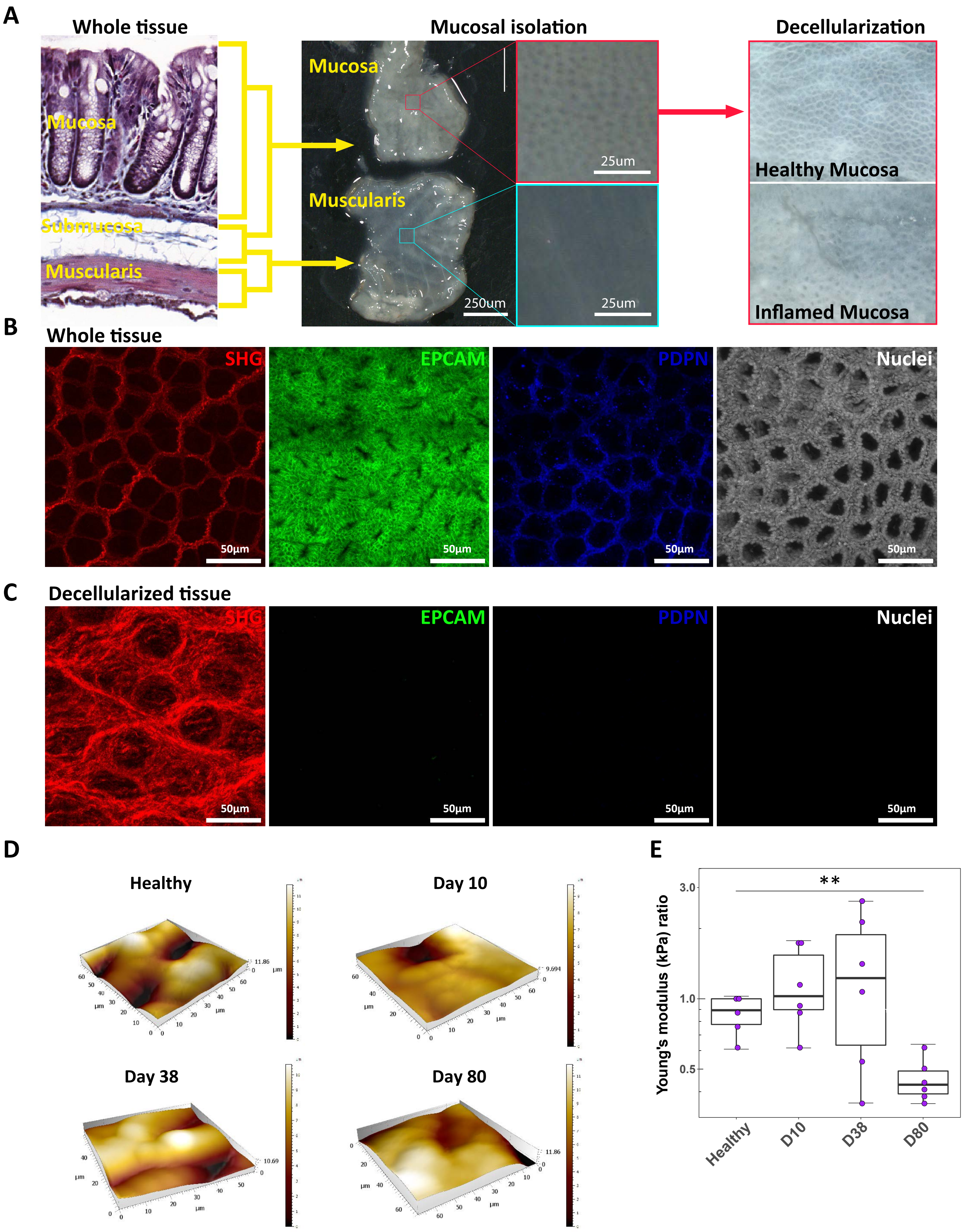
A
Whole tissue
Mucosa
Submucosa
Muscularis
Mucosal isolation
Mucosa
Muscularis
250um
25um
25um
Decellularization
Healthy Mucosa
Inflamed Mucosa
B
Whole tissue
SHG
EPCAM
PDPN
Nuclei
50µm
C
Decellularized tissue
SHG
EPCAM
PDPN
Nuclei
50µm
D
Healthy
Day 10
Day 38
Day 80
µm
E
Young's modulus (kPa) ratio
3.0
1.0
0.5
**
Healthy
D10
D38
D80

Figure S5.

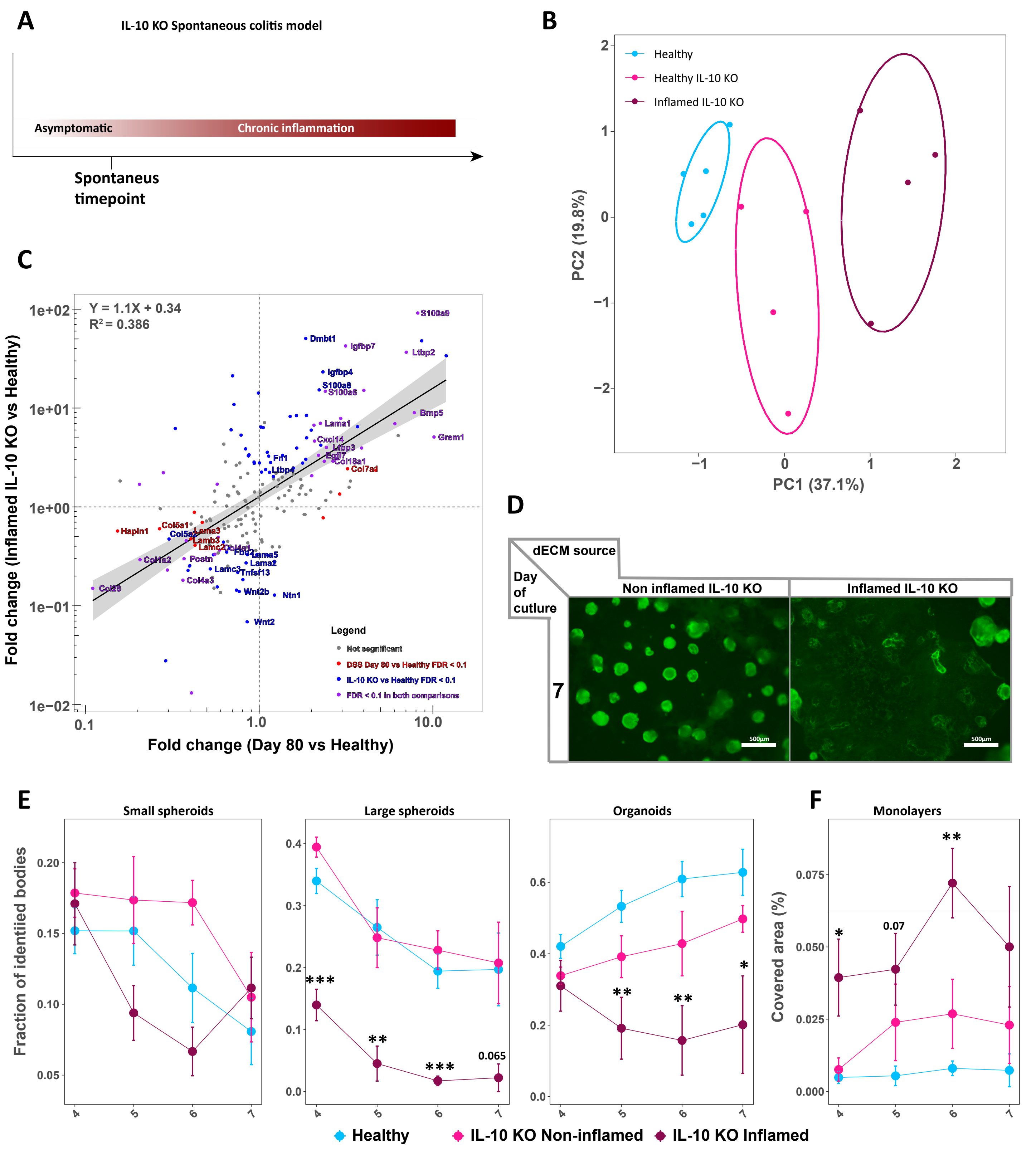

A
IL-10 KO Spontaneous colitis model
Asymptomatic
Chronic inflammation
Spontaneus timepoint
B
Healthy
Healthy IL-10 KO
Inflamed IL-10 KO
PC2 (19.8%)
PC1 (37.1%)
C
Y = 1.1X + 0.34
R² = 0.386
Fold change (Inflamed IL-10 KO vs Healthy)
Fold change (Day 80 vs Healthy)
Legend
Not segnificant
DSS Day 80 vs Healthy FDR < 0.1
IL-10 KO vs Healthy FDR < 0.1
FDR < 0.1 in both comparisons
D
dECM source
Day of cutlure
Non inflamed IL-10 KO
Inflamed IL-10 KO
7
E
Small spheroids
Large spheroids
Organoids
Fraction of identiled bodies
F
Monolayers
Covered area (%)
Healthy
IL-10 KO Non-inflamed
IL-10 KO Inflamed

Figure S6.

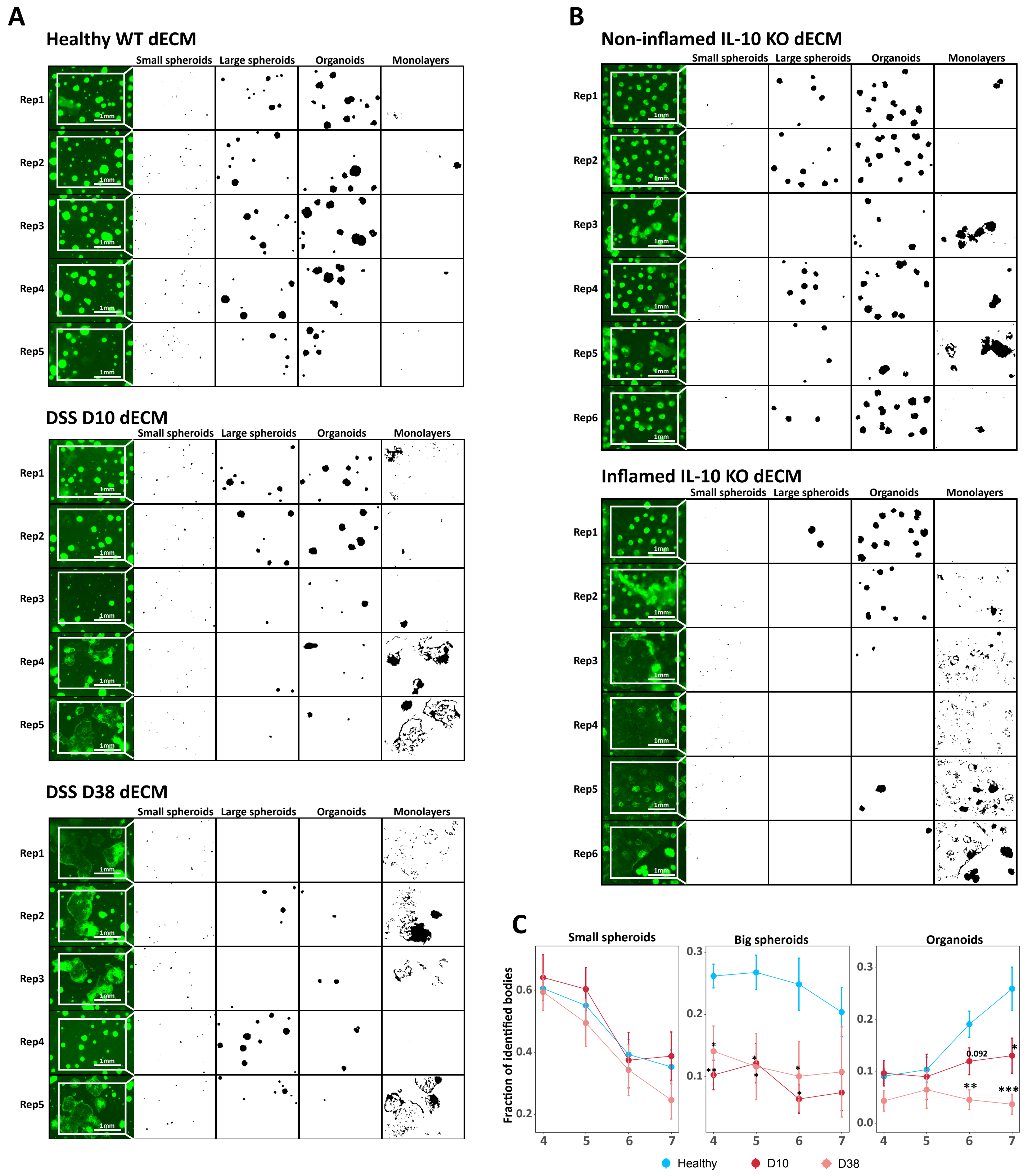
A
Healthy WT dECM
Small spheroids
Large spheroids
Organoids
Monolayers
Rep1
Rep2
Rep3
Rep4
Rep5
1mm
DSS D10 dECM
DSS D38 dECM
B
Non-inflamed IL-10 KO dECM
Rep6
Inflamed IL-10 KO dECM
C
Small spheroids
Big spheroids
Organoids
Fraction of identified bodies
0.092
Healthy
D10
D38

Figure S7.

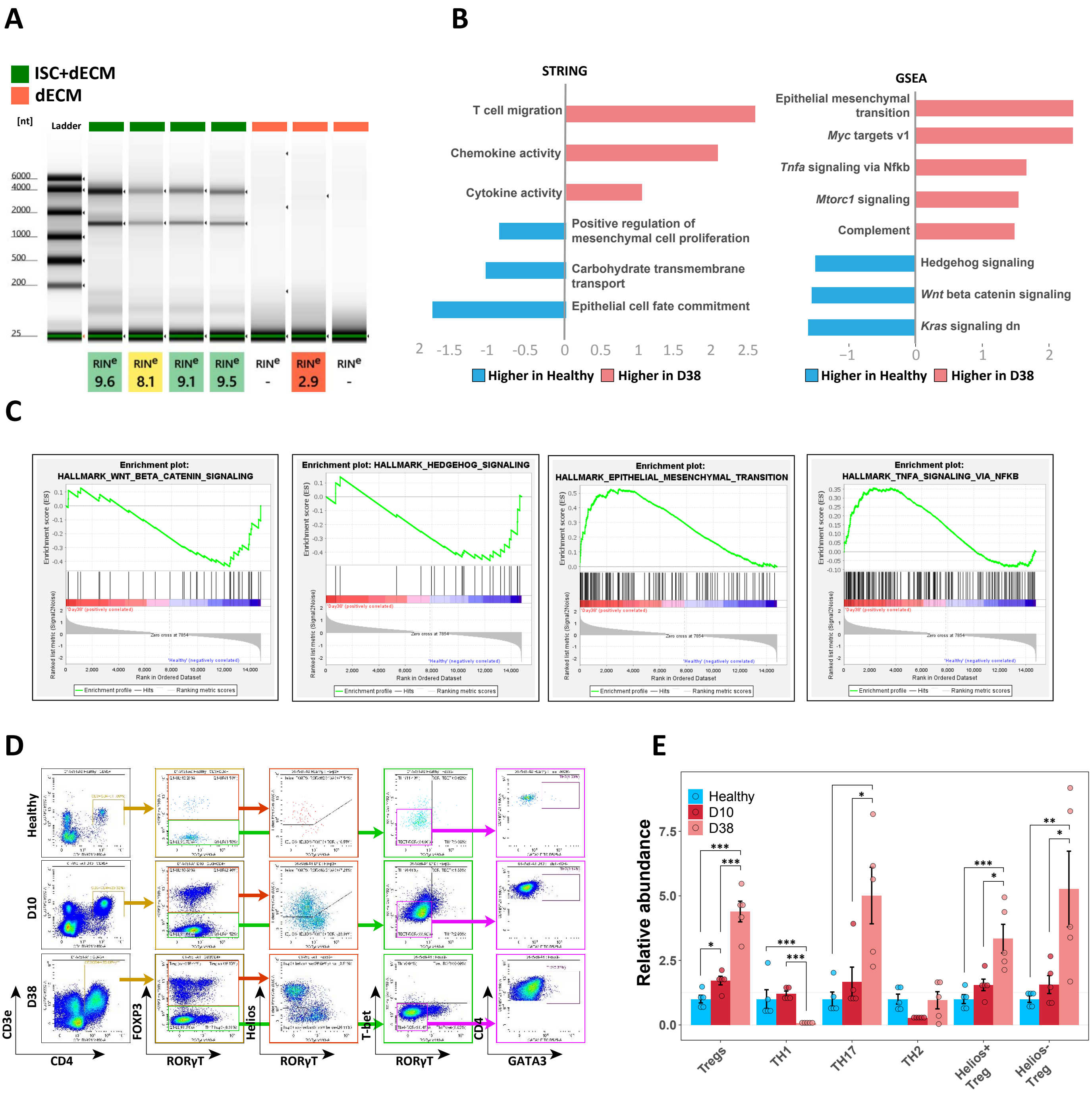

A
ISC+dECM
dECM
[nt]
Ladder
6000
4000
2000
1000
500
200
25
RINe 9.6
RINe 8.1
RINe 9.1
RINe 9.5
RINe -
RINe 2.9
RINe -
B
STRING
T cell migration
Chemokine activity
Cytokine activity
Positive regulation of mesenchymal cell proliferation
Carbohydrate transmembrane transport
Epithelial cell fate commitment
Higher in Healthy
Higher in D38
GSEA
Epithelial mesenchymal transition
Myc targets v1
Tnfa signaling via Nfkb
Mtorc1 signaling
Complement
Hedgehog signaling
Wnt beta catenin signaling
Kras signaling dn
Higher in Healthy
Higher in D38
C
Enrichment plot: HALLMARK_WNT_BETA_CATENIN_SIGNALING
Enrichment plot: HALLMARK_HEDGEHOG_SIGNALING
Enrichment plot: HALLMARK_EPITHELIAL_MESENCHYMAL_TRANSITION
Enrichment plot: HALLMARK_TNFA_SIGNALING_VIA_NFKB
Enrichment score (ES)
Ranked list metric (Signal2Noise)
Rank in Ordered Dataset
Enrichment profile
Hits
Ranking metric scores
D
Healthy
D10
D38
CD3e
CD4
FOXP3
RORγT
Helios
T-bet
GATA3
E
Healthy
D10
D38
Relative abundance
Tregs
TH1
TH17
TH2
Helios+ Treg
Helios- Treg
